# From cellular stress to cytokine modulation: An *in vitro* study of intestinal effects of major *Alternaria* mycotoxins

**DOI:** 10.64898/2026.10.30.761907

**Authors:** Anita Solhaug, Lada Ivanova, Vanessa Partsch, Rugile Jakaviciute, Gunnar Sundstøl Eriksen

## Abstract

Mycotoxins are frequently detected in food, making the gastrointestinal tract the first organ exposed. Here, we used various *in vitro* intestinal cell models to explore the barrier and immune responses to the Alternaria mycotoxins alternariol (AOH), alternariol methyl ether (AME), altenuene (ALT), altertoxin I (ATX-I), tentoxin (TEN), and tenuazonic acid (TeA). The mycotoxins exhibited low toxicity in undifferentiated Caco-2 cells, but AOH, AME, and ATX-I affected the cell cycle and reactive oxygen species (ROS) levels. None of the toxins alone or in combination with IL-1β had any effect on the barrier integrity or viability of the differentiated Caco-2/HT29-MTX-E12 co-culture. Furthermore, AOH and AME reduced IL-1β-induced IL-8 secretion, whereas TeA increased the secretion, demonstrating immunomodulatory effects. For verification, we used the more *in vivo*-like EpiIntestinal™ model, which is based on human primary cells. AOH, AME, and TeA had minor effects on viability without compromising barrier integrity; also, cytokine secretion was less affected than in the co-culture model. Studies on uptake confirmed low absorption of AOH, AME, and TeA. Overall, the studies indicate a low absorption and moderate effects of the toxins on the gastrointestinal tract.

## 1. Introduction

*Alternaria* fungi are widespread soil-living fungi. They are both plant pathogens and saprophytes and infect a wide range of plants, including crop plants. Some *Alternaria* species and their toxins are specific to certain host plants, while others grow on a wide range of plants (EFSA 2011). *Alternaria* seems to have a remarkable ability to adapt to the environmental conditions and is consequently found in both humid and semi-dry regions. *Alternaria* fungi produce a wide range of toxic and potentially toxic secondary metabolites (EFSA 2011), which may pose a risk to human and animal health. The mycotoxins produced by the *Alternaria* fungi are ubiquitous in the environment. Toxin production may also occur at lower temperatures, so food refrigerated during transportation and storage may also be contaminated (EFSA 2011).

Occurrence data for Alternaria mycotoxins in food are mostly available for alternariol (AOH), alternariol monomethyl ether (AME), tenuazonic acid (TeA), tentoxin (TEN), and altenuene (ALT), while very little information is currently available for other Alternaria toxins, including the perylene quinones, e.g. altertoxin I, II, and III. (ATX-I, -II, and -III). Currently, there are no regulations on Alternaria toxins in food or feed. Although human dietary exposure is estimated to be low, it exceeds the threshold of toxicological concern for potential genotoxic compounds (EFSA 2011). There is a lack of toxicological data on these mycotoxins, and *in vitro* studies may therefore help to characterise the hazards and clarify their mechanisms of action (Snapkow et al. 2024). Alternaria toxins have frequently been reported to induce cytotoxicity in various cell lines, with highly variable IC_50_ values across compounds. While the perylene quinone, ATX-II, is found to be cytotoxic in the low micromolar range (0.25 – 16 µM), considerably higher concentrations are needed for AOH and AME (3-120 µM) and TeA (70 – 140 µM), while. TEN was non-cytotoxic in the cell lines tested. The underlying mechanism of cytotoxicity induced by AOH, AME, and the perylene quinones is thought to involve interference with topoisomerase II, leading to DNA damage. ATX-II appears to be even more genotoxic than AOH, possibly by inducing DNA adducts through its epoxide-bearing perylene quinone structure in addition to the interaction with topoisomerase II (Fleck et al. 2012; Louro et al. 2024). Only very few studies on genotoxicity of TeA have been conducted TEN, and ALT, but so far, they have not been found to be genotoxic (Louro et al. 2024). The major mechanism of action of TeA is believed to be inhibition of protein synthesis (Kumari and Tirkey 2019). Furthermore, several Alternaria mycotoxins, including AOH, AME, TEN, ALT, ATX-I and ATX-II are reported to be immunosuppressive by inhibiting proinflammatory responses in human epithelial cells, human colon carcinoma cells, as well as in murine and human monocytes/macrophages (Aichinger et al. 2021; Crudo et al. 2025; Del Favero et al. 2020; Kollarova et al. 2018; Partsch et al. 2025; Partsch et al. 2026; Solhaug et al. 2015). The main mechanism is thought to involve interference with NF-κB activation, leading to decreased secretion of proinflammatory cytokines. Such immunosuppressive effects might increase susceptibility to certain diseases among people with chronic exposure to these mycotoxins.

The epithelium of the gastrointestinal tract (GIT) serves as a vital barrier against foodborne xenobiotics and is the primary site for nutrient absorption. Therefore, GIT cell models are essential *in vitro* systems for studying Alternaria toxin uptake and toxicity. Caco-2 cells, derived from a human colorectal carcinoma, are a well-characterized cell culture system. Proliferating Caco-2 cells may represent proliferative intestinal crypt cells, while fully differentiated Caco-2 monolayers that, display mature enterocyte-like morphology and function, comparable to those of the small intestine. Caco-2 cells cultured together with mucus-producing goblet cells (e.g. HT29-MTX-E12 has been proposed as a more physiologically relevant model as it forms an intestinal barrier with more *in vivo*-like TEER values and active mucus production. However, this model still has serious drawbacks as it has weak expression of several metabolic enzymes, such as CYP3A4, and variable expression of several transporters (Fedi et al. 2021; Markus et al. 2021). To overcome this, a human primary cell-based organotypic small intestine tissue model, the EpiIntestinal™ tissues provided from MatTek Life Sciences can be used. These tissues exhibit well-polarised morphology, form villi, brush borders, and tight junctions, and express efflux transporters and drug-metabolising enzymes that are more similar to those in *in vivo* intestinal tissues, making them a better model for human *in vivo* exposure (Markus et al. 2021).

Here, we have investigated the effects of six major Alternaria mycotoxins: AOH, AME, TeA, ATX-I, TEN, and ALT, on *in vitro* intestinal cell models, aiming to clarify whether they are cytotoxic and can modulate gastrointestinal inflammatory responses. First, we employed undifferentiated Caco-2 cells in the initial screening experiments to evaluate cytotoxicity, cell cycle progression, and reactive oxygen species (ROS) production. To assess effects on the barrier, mucus- and cytokine production, we used the differentiated co-culture of Caco-2 and HT29-MTX-E12 cells. Lastly, we used the more human-*in vivo*-like EpiIntestinal™ tissues to verify our key findings and to measure uptake of the most abundant mycotoxins: AOH, AME, and TeA.

## 2. Materials and methods

An overview of the methods, toxin concentrations, and exposure times used across the different cell models is provided in **Supplementary Table S1**.

### 2.1 Toxins and exposure

AOH and AME were synthesised by F. Reith, L Oltmanns, V. Prisyazhnoy and RD Süssmuth (Technical University of Berlin, Germany). AOH was synthesised according to Won (2015). The convergent synthesis featured a formylated aryl bromide and a desymmetrised aryl borane. Both were connected during a Pd-mediated Suzuki coupling. The final ring closure of the oxidised biaryl was performed with an ester formation during demethylation of all remaining alcohols. The product was isolated using reverse phase column chromatography and the identity of the material was confirmed by analytical HPLC-ESI-MS(MS) and ^1^HNMR spectroscopy. The convergent synthesis of AME was carried out according to Mikula et al. (Mikula et al. 2013). The building blocks were prepared from orcinol and 3,5-dimethoxyaniline, respectively and coupled using diethylchlorophosphate (DECP). The benzocoumarin core structure was established by intramolecular cyclisation of the iodoresorcylic acid phenyl ester. Selective demethylation of alternariol 2,4-dimethyl ether (ADME) with AlCl_3_/NaI yielded AME which was isolated using reverse-phase column chromatography. The identity of the material was confirmed by analytical HPLC-ESI-MS(MS) and ^1^HNMR spectroscopy. TeA (#202357) was purchased from Santa Cruz Biotechnology (TX, USA). ALT, (#21130), ATX-I, (#21129) and TEN, (#11442) were purchased from Cayman Chemical (MI, USA). The toxins were dissolved in DMSO (Sigma-Aldrich, Burlington, MA, USA) and stored in amber glass vials at -20 °C in 20 µL aliquots. The highest toxin concentrations used in the experiments are primarily determined by the toxin’s solubility and the need to avoid excessive DMSO concentrations during exposure (Solhaug et al. 2025). Upon exposure, the cell culture medium was replaced with medium containing the toxin. For exposure to a combination of toxin and IL-1β, the cells were first incubated with toxins for 2 h then IL-1β (0.5 ng/mL, Invivogen, CA, USA) was added apically. The final DMSO concentration in the cell culture was 0.2%, except in the first dose-response experiment (**Fig. 1**), where it was 0.4% due to the high toxin concentrations tested. All exposures within the same experiment had similar DMSO concentrations. DMSO had no effect on the parameters measured.

**Fig. 1:**
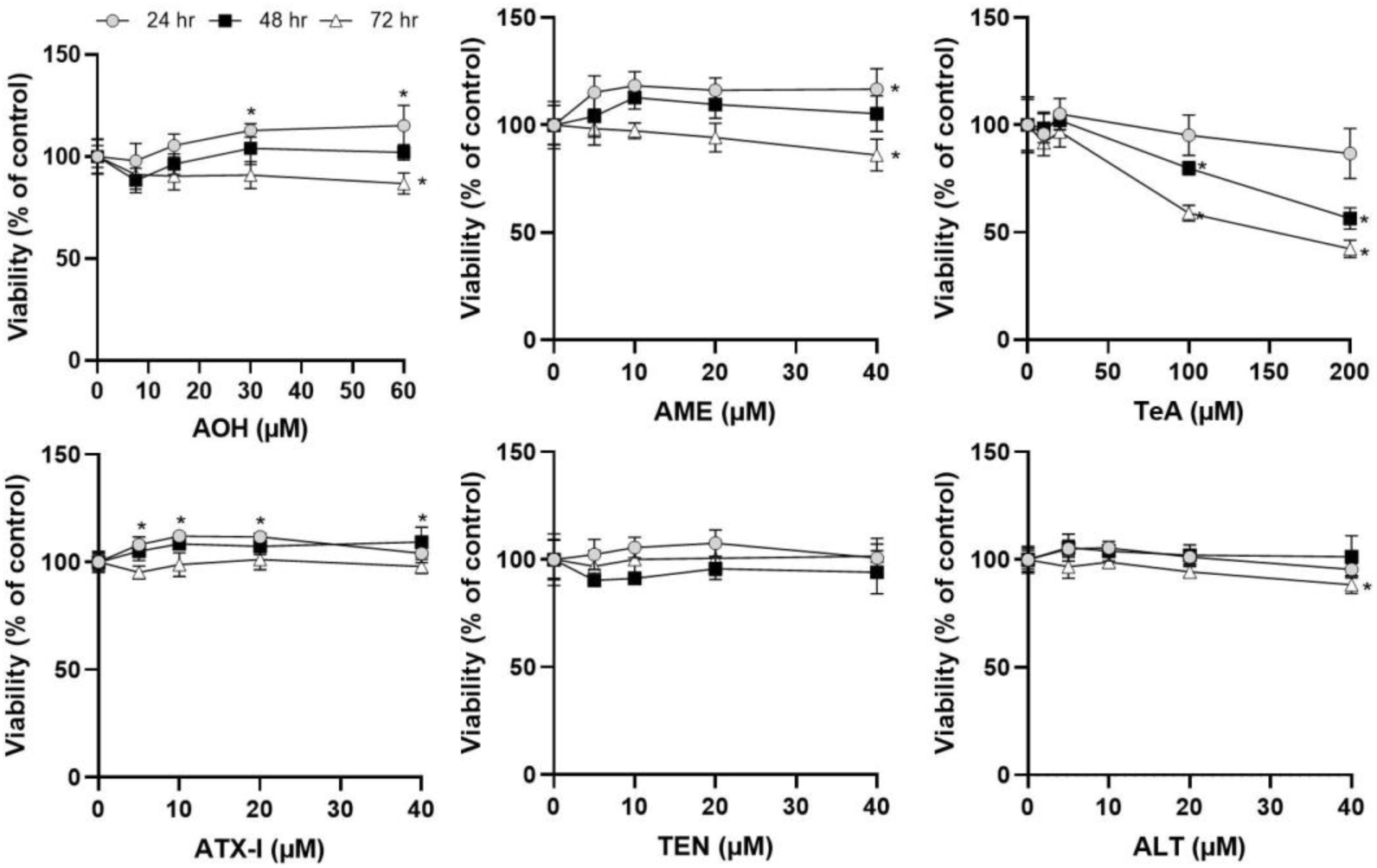
Undifferentiated Caco-2 cells were treated with Alternaria mycotoxins at the concentrations and times as indicated, and viability / metabolic activity was measured by the Alamar Blue assay. The data are expressed as the mean ± SD from three to six independent experiments, each performed in triplicate. Significant difference (p< 0.05) compared to control (0 µM) is indicated with an asterisk (*). One-way ANOVA with Dunnett’s post hoc test.

### 2.2 Cell cultivation

The Caco-2 (# 09042001) and HT29-MTX-E12 (#12040401) cell lines were obtained from European Collection of Authenticated Cell Cultures (ECACC). The cells were cultivated in 72 cm^2^ vented flasks with Dulbecco’s Modified Eagle medium (DMEM glutamax, Gibco, Thermo Fisher, MA, USA), supplemented with 1% non-essential amino acids (NEAA, Gibco, Thermo Fisher), 1% penicillin-streptomycin (Gibco, Thermo Fisher), and 10% heat-inactivated fetal bovine serum (FBS, USA origin, Gibco, Thermo Fisher) at 37 °C, 5% CO_2_. The cell lines were routinely screened for mycoplasma contamination using the MycoAlert® Mycoplasma Detection Kit (Lonza, Basel, Switzerland). Subcultivation was performed twice a week, when the culture reached approximately 70% confluence, using TrypLE (Gibco, Thermo Fisher). <u>Caco-2 monoculture:</u> Cells were plated in standard cell culture plates at a density of 1.3 × 10^5^ cells/cm^2^ the day before exposure. For the uptake studies, Caco-2 cells were seeded into standing cell culture transwell inserts (Millicell® cell culture inserts, 0.4 µm, 0.6 cm^2^, PIHP01250, Sigma-Aldrich) at a density of 1 × 10^5^ cells/cm^2^. The cells were then grown for 21 days, with medium refreshment every Monday, Wednesday, and Friday. <u>Co-culture:</u> A co-culture consisting of Caco-2 and HT29-MTX-E12 cells (9:1 ratio) was plated into hanging cell culture transwell inserts (Millicell ® cell culture inserts, 0.4 µm, PTHT24H48, Sigma-Aldrich) at a density of 1 × 10^5^ cells / cm^2^. The cells were then grown for 21 days, with medium refreshment every Monday, Wednesday, and Friday. <u>Primary human intestinal cells:</u> 3D EpiIntestinal™ tissues (standing inserts, 24-well format, 0.6 cm^2^) were obtained from MatTek Life Sciences (Sartorius, Bratislava, Slovak Republic) and cultured according to the manufacturer’s instructions using the provided cell culture medium. These tissue models are composed of normal, human-derived small intestinal cells. The EpiIntestinal™ tissue model (SMI-100) is a cell-based 3D model that incorporates enterocytes, Paneth cells, M cells, tuft cells, and intestinal stem cells into a highly differentiated, polarised epithelium. The EpiIntestinal™ FT tissue (SMI-200-FT) also includes an underlying lamina propria containing normal human intestinal fibroblasts, making it useful for studying inflammation. The SMI-200-FT model was used for barrier, viability, and cytokine secretion experiments, while the SMI-100 was used for uptake studies.

### 2.3 Gut-On-Chip model

A co-culture of Caco-2 and HT29-MTX-E12 cells (9:1) was plated onto hanging inserts as described in section 2.2. After 21 days of differentiation, the inserts were transferred into single MIVO single-flow chambers (React4Life, Genova, Italy). The basolateral flow rate of 2.3 mL/min, corresponding to physiological velocities (personal communication, React4Life), was achieved using a peristaltic pump in a closed circuit. A static control was also included.

### 2.4 Viability / metabolic activity

The metabolic activity / viability of the cells was measured using the Alamar Blue assay according to the manufacturer’s protocol (Invitrogen, Thermo Fisher). The dark blue oxidised form of Alamar Blue (resazurin) is reduced to a highly fluorescent form (resorufin) by mitochondrial and cytoplasmic enzymes (Solhaug et al. 2025). The measured fluorescence intensity is thus proportional to the number of viable cells. The fluorescence, Excitation (Ex) 555 nm / Emission (Em) 585 nm, was quantified using Spectramax i3x plate reader (Molecular Devices, San Jose, CA, USA).

### 2.5 Cell cycle

For cell cycle analysis, the Caco-2 cells were harvested, washed once with PBS (Gibco, Thermo Fisher), and fixed with ice-cold 70% ethanol overnight at –20 °C. The cells were then washed with PBS, incubated with 10 µg/mL propidium iodide (PI, Molecular probe, Thermo Fisher) / 100 µg/mL RNase A (Invitrogen, Thermo Fisher) in PBS for 30 min at 37 °C before analyses on a flow cytometer (Accuri C6, BD, Franklin Lakes, NJ, USA). Single cells were gated, and a minimum of 10,000 cells were acquired within the gate of interest and analysed using the FL2-A (585/40) filter.

### 2.6 ROS

ROS production was detected by using the oxidation-sensitive fluorescent probe, CM-H_2_DFDA (Molecular Probe, Invitrogen, Thermo Fisher). Since fluorescent probes are known to be good substrates for efflux pumps (Zorova et al. 2018), which are highly expressed in barrier cell types such as Caco-2 cells, the incubation of CM-H_2_DFDA was performed in the presence of the efflux-pump inhibitor probenecid (Gollapudi et al. 1997). The cells were first pre-incubated with CM-H_2_DFDA (10 µM) in combination with probenecid (1 mM, water-soluble, #50027, Biotium, Fremont, CA, USA) in PBS for 30 min, followed by exposure to Alternaria mycotoxins in complete cell culture medium for 2 hr and 24 hr. The cells were then detached using TrypLE, resuspended in PBS, and analysed by flow cytometry (Accuri C6, BD, Franklin Lakes, NJ, USA) with the FL1-A filter (530/30 nm filter).

### 2.7 Barrier

Barrier integrity was assessed by measuring transepithelial electrical resistance (TEER) using a Millicell RES Voltmeter (Millipore, Burlington, MA, USA).

### 2.8 Cytokines

<u>ELISA:</u> After exposure, IL-8 and IL-6 levels were measured by enzyme-linked immunosorbent assay (ELISA), according to the manufacturers’ guidelines (CHC1303 and CHC1263, respectively; Life Technologies, Invitrogen, Thermo Fisher). The absorbance was measured using a plate reader (Spectramax). <u>Cytokine bead assay (CBA):</u> After exposure, the harvested cell culture medium was simultaneously studied for secreted cytokines (IL-1β, TNF, IL-6, IL-8, IL-10, IL-12p70) using the BD Cytometric bead Array (CBA) Human inflammatory cytokines kit (BD Biosciences, # 551811) and analysed by flow cytometry (Accuri C6).

### 2.9 MUC5AC secretion

Detection of secreted MUC5AC was performed using a protocol modified from Wan et al. (2014). The apical cell culture medium was diluted 1:4 in cell culture medium and mixed 1:2 with coating buffer (Sodium carbonate 0.16 M/ hydrogen Carbonate 0.034 M; pH 9.6) into an ELISA plate (Maxisorp, Thermo Fisher) and evaporated to dryness at 40 °C overnight. The ELISA plate was then washed twice with PBS/Tween-20, 0.05% (PBST), followed by blocking with assay buffer (PBST containing 2% BSA (Sigma)). After 1 h, the plate was washed three times with PBST and incubated with MUC5AC antibody (1:1000, #MA1-21907, Thermo Fisher) diluted in assay buffer for 2 hr on a plate shaker (700 rpm). The plate was washed three times with PBST and then incubated with HRP-conjugated goat anti-mouse antibody (1:2000; #7076, Cell Signaling Technology, MA, USA). After 2 h, on a plate shaker (700 rpm), the plate was washed three times with PBST and the colour reaction was developed with 3,39,5,59-tetramethylbenzidine (TMB) peroxidase solution (Neogen, Lansing, MI, USA) and stopped with 10% H_2_SO_4_. Absorbance was read at 450 nm with a plate reader (Spectramax).

### 2.10 Transport studies

Differentiated Caco-2 cells or EpiIntestinal™ tissues (SMI-100), grown in standing cell culture transwell inserts, were placed in 12-well plates with 300 µL of cell culture medium in the apical compartment and 1500 µL in the basolateral compartment. Exposures were performed apically. Incubation aliquots (150 µL) collected in apical and basolateral sides at specific time points were mixed on ice with a 1:1 (v/v) ratio of ice-cold 100% acetonitrile (ACN). After vortexing and centrifugation at 20,000× g for 10 min at 4°C, the samples were stored at −20 °C until LC-HRMS/MS analysis. On the day of analysis, samples were thawed on ice and centrifuged again if precipitation was observed. Before UPLC–HRMS/MS analysis, samples were filtered using Thomson Single StEP nano filters. For intracellular toxin analysis, cells were first scraped with 100 µL lysisbuffer (CellLytic^TM^ ,Sigma-Aldrich) as previously done for intracellular Enniatin B1 analysis (Ivanova et al. 2010) to ensure efficient cell lysis and recovery of intracellular and membrane bound compounds, then homogenised using QIAshredder”. In contrast to direct lysis in ACN, this method also enables later analysis of proteins (e.g. western blotting) if needed. The cell lysis was then mixed with an equal volume of ice-cold 100% ACN (1:1, v/v), for protein precipitation and centrifugation at 20,000 × g for 10 min at 4 °C. The resulting supernatants were filtered through Thomson Single StEP nano filters before UPLC–HRMS/MS analysis (Thomson Instrument Company, CA, USA).

### 2.11 Liquid chromatography-high-resolution mass spectrometry (LC-HRMS)

Alternaria toxins were determined using a Q Exactive™ hybrid quadrupole–Orbitrap mass spectrometer equipped with a heated electrospray ionisation source (HESI-II) and coupled to a Vanquish UHPLC system (Thermo Fisher Scientific). Chromatographic separation was performed on a Waters Acquity BEH C18 column (50 × 2.1 mm, 1.7 μm; Waters GmbH, Eschborn, Germany) in combination with an Acquity BEH C18 VanGuard^TM^ pre-column (2.1 × 5 mm, 1.7 µm), maintained at 40 °C. The sample injection volume was 1 μL. The mobile phase consisted of 1 mM aqueous ammonium bicarbonate (NH_4_HCO_3_) and methanol (MeOH) at a flow rate of 0.4 mL/min. The gradient started at 2% MeOH, increased to 10% at 2 min, held until 2.3 min, and then increased to 95% MeOH at 6.5 min. The total runtime was 10 min, including a 1 min cleaning step at 100 % MeOH and a 1 min equilibration. The HESI-II source was operated in negative-ion mode with a spray voltage of 3.2 kV, capillary temperature of 280 °C, source temperature of 300 °C, sheath gas flow of 35 L/min, auxiliary gas flow of 10 L/min, and S-lens RF of 55. Full-scan mass spectra were acquired at 70,000 FWHM (*m/z* 150–600) with an AGC target of 1 × 10⁶ and a maximum injection time of 250 ms, using ddMS²-top5 and an inclusion list (**Table 1**). HRMS/MS spectra were recorded at 17,500 FWHM with HCD and stepped normalised collision energies of 20, 35, and 60 eV, with AGC target 5 × 10⁴, maximum injection time 150 ms, and an isolation window of *m/z* 1.4. The instrument was externally calibrated once per week in positive and negative ion modes using a Pierce® calibration solution (Thermo Fisher Scientific) according to the manufacturer’s recommendations. Data were processed using Xcalibur 4.2.28.14. Identification of Alternaria toxins was based on retention time and HRMS/MS fragment comparison with reference standards, with a mass accuracy tolerance of ≤ 5 ppm. Quantitative analysis was performed using external calibration with matrix-matched standards over a concentration range of 1–200 ng/mL. Matrix-matched calibration standards were prepared by diluting the culture medium (1:1, v/v) with 100% ACN, which was also used to dilute samples that exceeded the calibration range. The method for determining Alternaria toxins was partially validated, and key performance parameters are summarized in **Table 1**. Calibration curves were prepared using standards in 50% ACN and in medium. Limits of detection (LOD) and quantification (LOQ) were calculated as LOD = 3SD/s and LOQ = 10SD/s, based on the standard deviation of the y-intercept and the slope of the average regression line based on the two calibration sets analysed on the same day (Shrivastava and Gupta 2011).

**Table 1.** UHPLC-HRMS parameters for the determination of AOH, AME and TeA and performance parameters including limits of detection and quantification (LOD and LOQ) in solvent and matrix-matched calibration, and matrix-induced signal suppression/enhancement (SSE).

| ***Compound*** | ***Ionisation mode<sup>a</sup>*** | ***Elemental formula*** | ***RT, min*** | ***Ion mass* (m/z)** | **LOD (ng/mL)** |  | **LOQ (ng/mL)** |  | **SSE%** |
|---|---|---|---|---|---|---|---|---|---|
|  |  |  |  |  | ***Pure std.*** | ***"Matrix"*** | ***Pure std.*** | ***"Matrix"*** |  |
| TeA | ES neg | C<sub>10</sub>H<sub>15</sub>NO<sub>3</sub> | 2.0 | 196.0979 | 0.9 | 3 | 2.9 | 8.1 | 119 |
| AOH | ES neg | C<sub>14</sub>H<sub>10</sub>O<sub>5</sub> | 5.5 | 257.0456 | 5.0 | 2 | 15 | 7.2 | 96 |
| AME | ES neg | C<sub>15</sub>H<sub>12</sub>O<sub>5</sub> | 6.4 | 271.0612 | 13 | 11 | 38 | 32 | 122 |

<sup>a</sup>[M-H]<sup>-</sup>

### 2.12 Calcein accumulation assay

Calcein-AM is a non-fluorescent, cell-permeable compound. Once inside the cell, it is hydrolysed by intracellular esterases to calcein, a highly fluorescent, hydrophilic molecule that is trapped within the cytoplasm. In cells overexpressing P-glycoprotein (P-gp), Calcein-AM is actively pumped out of the cell before it can be converted, resulting in low intracellular fluorescence. P-gp inhibitors block this efflux, leading to the accumulation of calcein and a corresponding increase in fluorescence intensity, which is directly proportional to the inhibitor’s potency (Tiberghien and Loor 1996). The cells were plated out in black 96-well plates (1 × 10^5^ cells / cm^2^) the day before the experiment. The cells were pre-incubated with Alternaria toxins or the efflux transporter P-gp inhibitor PSC833 (5 µM; Sigma-Aldrich) as a positive control, for 5 min before the addition of Calcein-AM (Sigma-Aldrich). After incubation, the medium was removed, and the cells were lysed with PBS containing 0.3% Triton. The cellular Calcein-AM uptake was then measured and quantified using a Spectramax i3x plate reader in fluorescence mode (Ex 494 nm / Em 517 nm). To ensure a representative readout of the fluorescent adherent cells, 37 points were acquired in each well using the plate reader’s well-scan function. After the initial optimisation experiments (**Supplementary Fig. S8**), 1 h incubation with 0.5 µM Calcein-AM was used in the study.

### 2.13 Statistical analysis

The data analyses were performed using GraphPad Prism (version 9.0.1.151). Statistical significance was considered at p < 0.05, and analysed using one- or two-way ANOVA with Dunnett’s post hoc test, as specified in the figure legends. In general, the number of independent experiments exceeds three. However, the cost of EpiIntestinal™ tissues is high, which limits our ability to use them; therefore, they are only used as two or three independent incubations. Nevertheless, most experiments are conducted across several model systems using methods that complement each other.

## 3. Results

### 3.1. Influence of Alternaria mycotoxins on cell growth and oxidative stress

Proliferating Caco-2 cells are often considered a model of intestinal crypt cells. Also, it is less cost- and time-consuming model to work with, as the cells grow on ordinary plastic plates and do not require 3 weeks of differentiation on transwell inserts, which also makes the model a good tool for initial toxicity screening. So, to explore the overall toxicity of six Alternaria toxins to intestinal cells, the simplest model in this study, undifferentiated Caco-2 cells, was exposed to the toxins over a wide concentration range. The highest toxin concentrations used are primarily determined by the toxin’s solubility and the need to avoid excessive DMSO concentrations during exposure (Solhaug et al. 2025). For TeA, higher concentrations were used than for the other toxins, due to its higher solubility and overall higher occurrence. The Caco-2 cells were treated with AOH (7.5, 15, 30, 60 µM), AME (5, 10, 20, 40 µM), TeA (10, 20, 100, 200 µM), ATX-I (5, 10, 20, 40 µM), TEN (5, 10, 20,

40 µM), and ALT (5, 10, 20, 40 µM) for 24, 48 or 72 h, and the viability, measured by metabolic activity; the Alamar Blue assay (**Fig. 1**). After 24 h exposure, AOH (30, 60 µM), AME (40 µM) and ATX-I (40 µM) increased metabolic activity (probably stress), while reduced metabolic activity was observed for AOH (60 µM), AME (40 µM), ALT (40 µM) after 72 h exposure, and for TeA (100 and 200 µM) after 48 and 72 h exposure. Detachment or apoptotic/necrotic morphology of the cells was not observed following exposure to any of the toxins, even after 72 h of exposure at the highest concentrations (**Fig. 2**), indicating that AOH, AME, TeA, and ALT reduce viability by affecting proliferation or overall metabolic activity rather than by causing cell death.

**Fig. 2:**
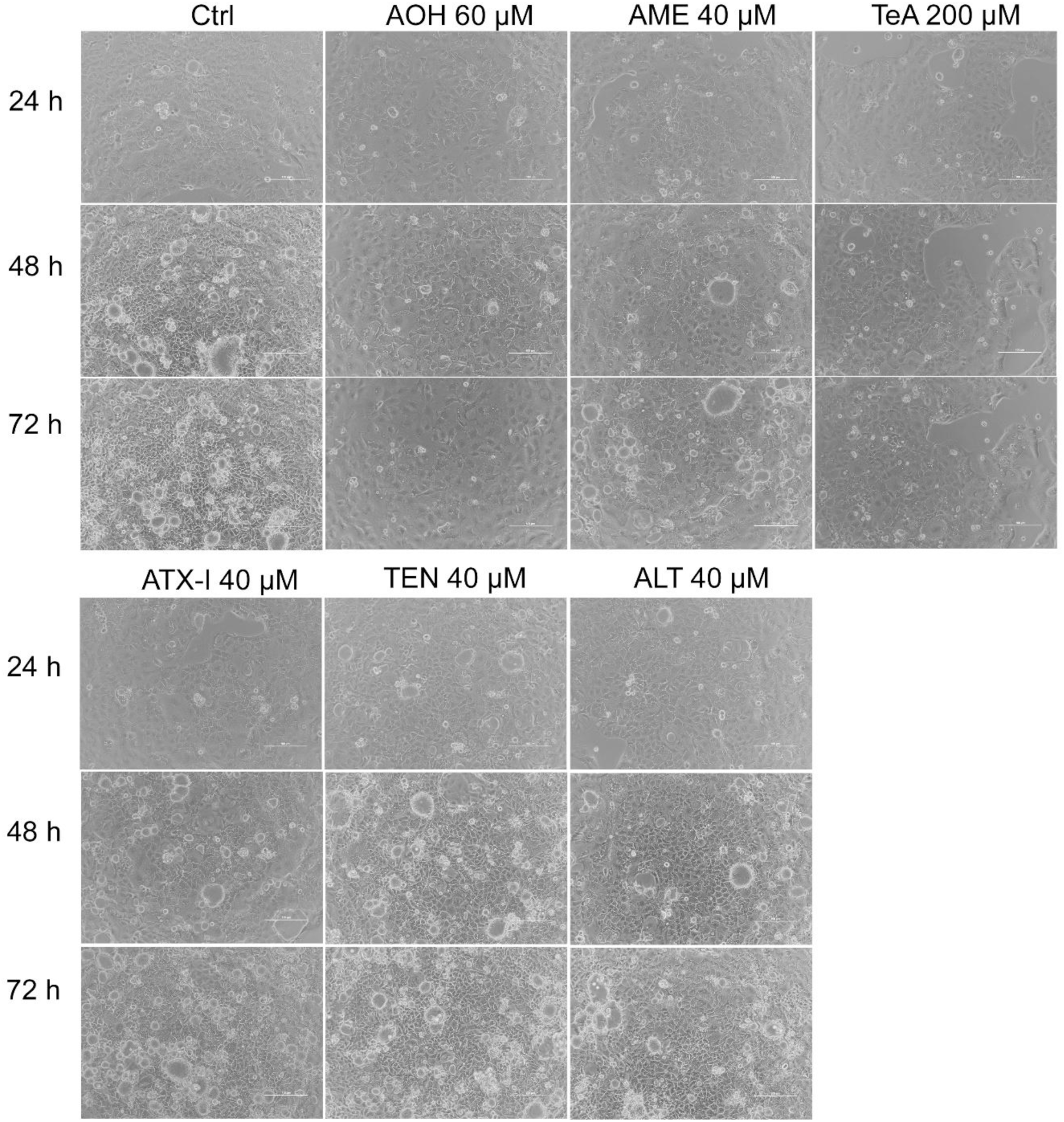
Undifferentiated Caco-2 cells were treated with Alternaria mycotoxins at the concentrations and times as indicated, and images were captured with a phase contrast microscope. Scalebar = 100 µm.

To further investigate toxin-mediated effects on cell growth, we analysed cell cycle progression. Sub-confluent Caco-2 cells exhibit a cell cycle duration of approximately 30 h; therefore, a 24-h exposure period was selected to enable detection of potential perturbations in cell cycle progression. Based on the Alamar Blue viability results (**Fig. 1**), the selected mycotoxin concentrations were non-cytotoxic and are considered relevant for more detailed *in vitro* studies, given the known occurrence data (EFSA 2011; EFSA 2016). Exposure to AOH, AME, and ATX-I resulted in a decrease in the proportion of cells in the G1 phase, accompanied by a corresponding increase in the fractions of cells in the S and G2 phases and a significant accumulation of cells specifically in the G2 phase was observed at the highest concentration of AOH and AME (**Fig. 3**). In contrast, TeA reduced cell viability, but had no effect on the cell cycle. No effect on the cell cycle was observed for TEN and ALT.

**Fig. 3:**
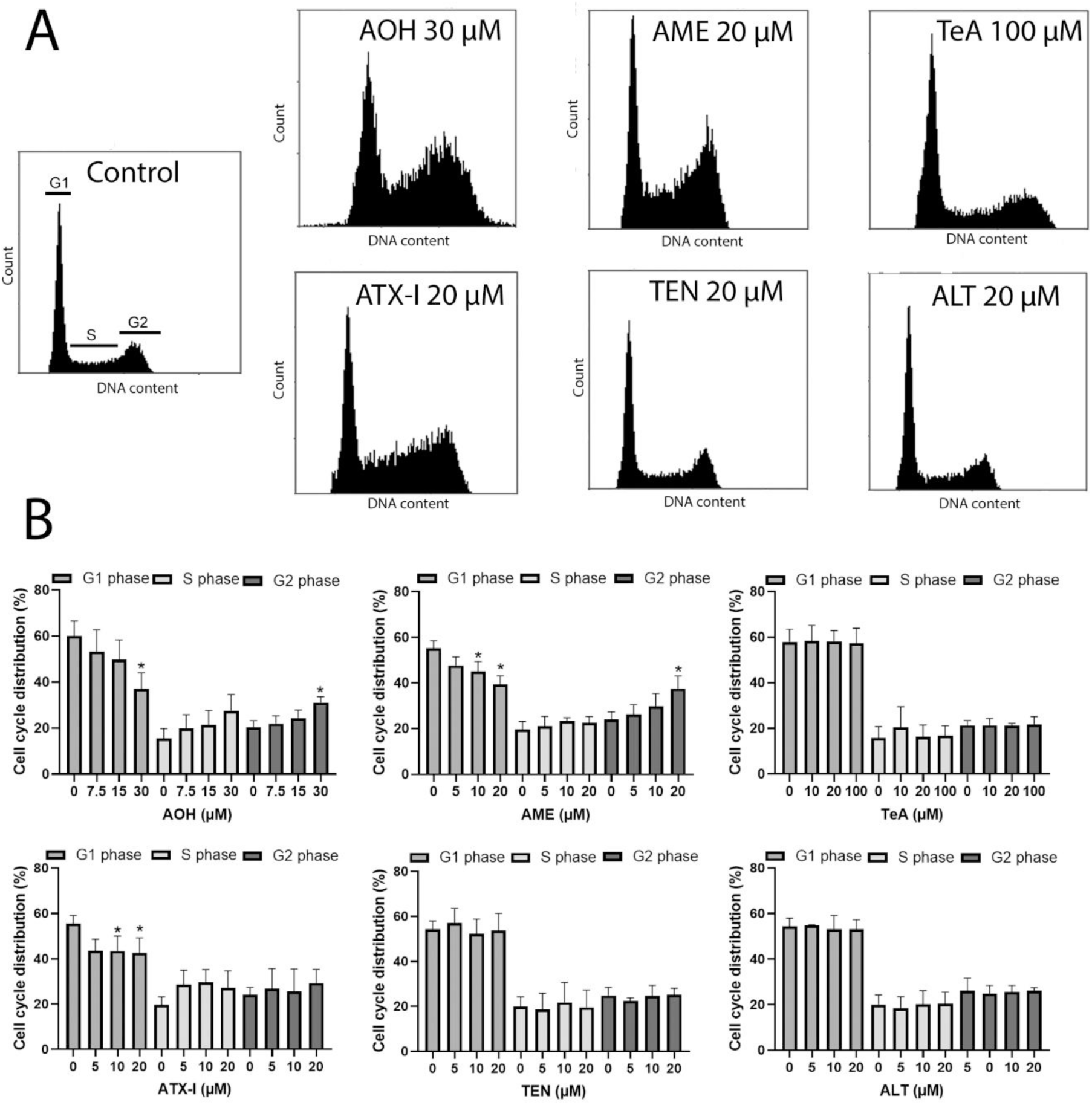
Undifferentiated Caco-2 cells were treated with Alternaria mycotoxins at the concentrations indicated for 24 h and analysed for cell cycle (**A**). The data (**B**) are expressed as mean + SD from four independent experiments. Significant difference (p< 0.05) compared to control (0 µM) is indicated with an asterisk (*). One-way ANOVA with Dunnett post hoc test.

Toxin exposure often leads to oxidative stress, with the production of ROS associated with biotransformation and/or mitochondrial dysfunction. Notably, AOH, AME, and ATX-I, which caused a reduction of cells in the G1 phase, also increased cellular ROS levels after 2 and 24 h exposure (**Fig. 4**).

**Fig. 4:**
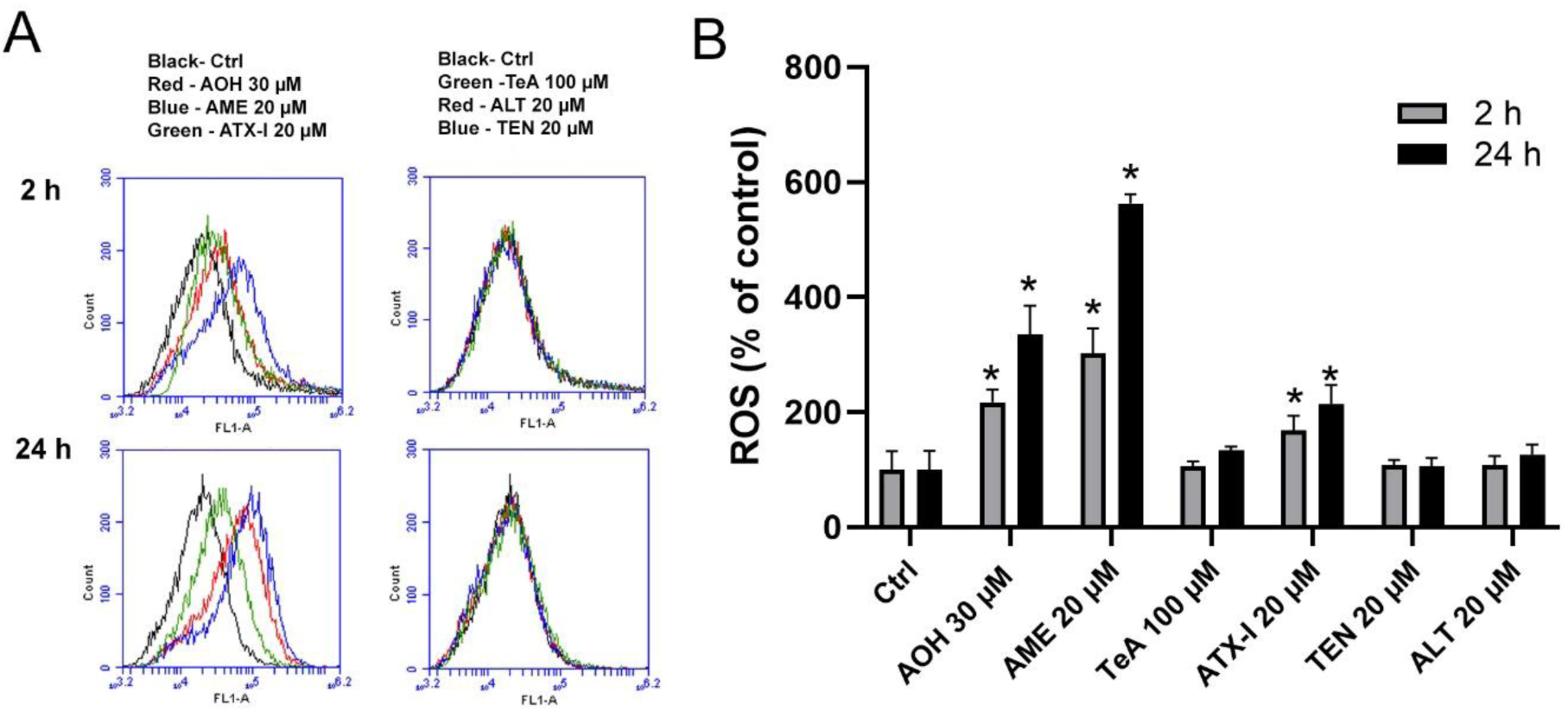
Undifferentiated Caco-2 cells were treated with Alternaria mycotoxins at the concentrations indicated for 2 or 24 h and analysed for ROS by flow cytometry. FL-1 represents green fluorescence; levels of ROS (**A**). The data are expressed as mean + SD from three to four independent experiments (**B**). Significant difference (p< 0.05) compared to its representative control (Ctrl) is indicated with an asterisk (*). Two-way ANOVA with Dunnett post hoc test.

### 3.2 Influence of Alternaria mycotoxins on intestinal barrier and inflammation

To test the effects of Alternaria toxins in a model system more closely resembling the intestine, we next used differentiated co-culture of Caco-2 and mucus-producing HT29-MTX-E12 cells. In contrast to undifferentiated Caco-2 cells, the co-culture is cultivated on transwell inserts, which allow differentiation of the monolayer. After 21 days, the co-culture has a stable TEER (**Supplementary, Fig S1**), which is an indicator of fully differentiated cells with a functional barrier (Fedi et al. 2021), and thus ready for the experiment. We focused on a non-cytotoxic concentration range and also included lower concentrations than used in the previous experiments to evaluate potential effects on barrier function and inflammatory responses at more realistic concentrations. To mimic an inflamed intestine, the toxins were also incubated with a low concentration of IL-1β, which induces low-level inflammation (Al-Sadi and Ma 2007). The exposure method was partly adapted from a study by Schmutz et al. (2019), which investigated the immunosuppressive potential of AOH in Caco-2 cells. Briefly, cells were either incubated with toxin alone (AOH 0.3, 3, 30 µM, AME, ATX-I, TEN, ALT 0.2, 2, 20 µM and TeA 2, 20, 100 µM) for 20 h, or cells were first pre-incubated for 2 hr with the toxin and then additionally stimulated with IL-1β (0.5 ng/mL) for a further 18 h. The concentration of IL-1β is selected to allow both inhibition of response and additional induction of response. The different parameters, viability (Alamar Blue), Barrier (TEER), IL-8 secretion, and MUC5AC secretion were measured from the same samples.

As with undifferentiated Caco-2 cells, the highest concentration of TeA reduced the metabolic activity, whereas the other toxins had no effect on the differentiated co-culture (**Fig. 5A**). Interestingly, even though metabolic activity decreased with TeA exposure, no effect of TeA was found on barrier integrity, assessed with TEER measurements (**Fig. 5B**). Likewise, none of the other toxins had any effects on the TEER. Altogether, these results further indicate that the toxins have low cytotoxicity and no functional effects on the tight junctions in the intestinal tract in the tested concentrations.

**Fig. 5:**
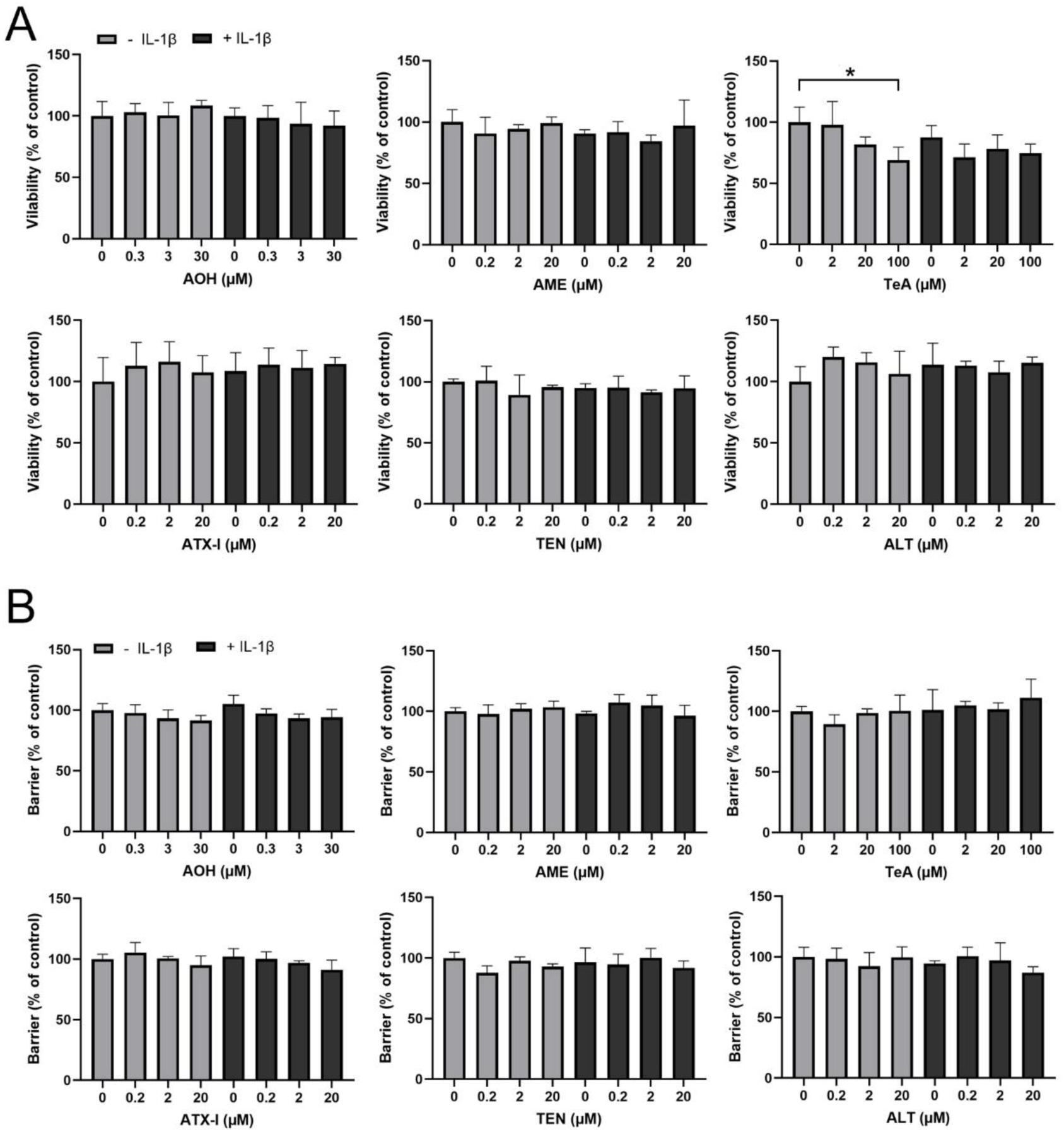
Differentiated Caco-2/HT29-MTX-E12 co-cultures were treated with Alternaria mycotoxins as indicated for 2 h before the addition of IL-1β (0.5 ng/mL) for additional 18 h, and viability (**A**) and Barrier integrity; TEER (**B**) was measured. The data are expressed as mean + SD from four to six independent experiments. Significant difference (p< 0.05) compared to ctrl (0 µM) is indicated with an asterisk (*). One-way ANOVA with Dunnett’s post hoc test.

IL-1β did not affect the viability or the barrier of the co-culture alone or in combination with the mycotoxins (**Fig. 5**). When we treated the cells with a low concentration of IL-1β (0.5 ng/mL), a moderate increase in IL-8 secretion (about 10-fold) was observed (**Fig. 6**). None of the toxins alone induced IL-8 secretion in the co-culture, but the highest tested concentrations of AOH (30 µM) and AME (20 µM) reduced IL-1β-induced IL-8. Interestingly, low AME concentration (0.2 µM) and the highest TeA concentration (100 µM) increased IL-1β-induced IL-8 secretion, indicating a potentiating effect on IL-1β induced inflammation.

**Fig. 6:**
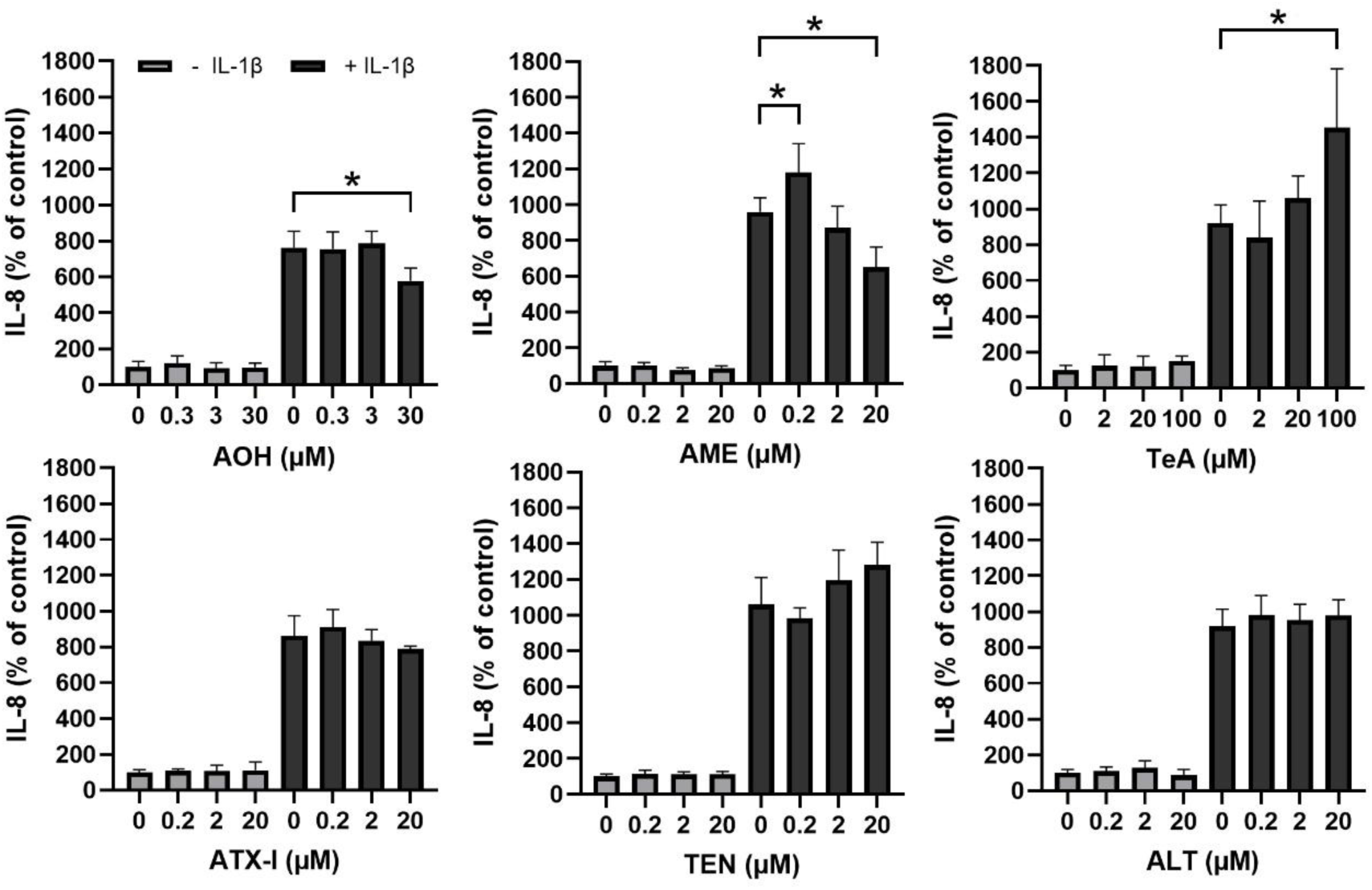
Differentiated Caco-2/HT29-MTX-E12 co-cultures were treated with Alternaria mycotoxins as indicated for 2 h before the addition of IL-1β (0.5 ng/mL) for additional 18 h, and IL-8 secretion was measured. The data are expressed as mean + SD from four to six independent experiments. Significant difference (p< 0.05) compared to ctrl (0 µM) is indicated with an asterisk (*). One-way ANOVA with Dunnett’s post hoc test.

To test the effects of physiological flow, we placed the co-culture in a gut-on-a-chip model, a single MIVO flow chamber with a basolateral flow rate of 2.3 mL/min, corresponding to physiological velocities. Regarding viability/metabolic activity (Alamar blue), AOH (30 µM), AME (20 µM) or TeA (100 µM) in combination with IL-1β had no effect on viability compared to IL-1β alone, with or without flow. However, a difference was observed between the static and the fluidic condition for AME + IL-1β, with somewhat lower viability when the cells were treated under fluidic condition (**Supplementary**, **Fig. S2A)**. Furthermore, it was a general tendency of lower IL-8 secretion under fluidic conditions, however a significant difference between the static and the fluidic condition was only found for TeA (100 µM) + IL-1β (**Supplementary, Fig S2B).** Notable, in contrast to our previous experiments (**Fig. 6**), AOH and AME only showed a tendency, but did not significantly reduce IL-1β induced IL-8 secretion under static conditions. It should, however, be noted that the higher variance observed in the static versus fluidic experiments **(Supplementary, Fig S2B)** compared with previous experiments under static conditions (**Fig. 6**) may have resulted in an underestimation of the effect. Lastly, we analysed MUC5AC as an indicator of mucin secretion from the HT29-MTX-E12 cells, using the static, co-culture model. Interestingly, low concentration of ALT (0.2 µM) increased MUC5AC secretion by 45% (**Supplementary, Fig. S3**), but no effect was observed in combination with IL-1β. None of the other mycotoxins tested had any effect, either alone or in combination with IL-1β.

To verify our findings from the Caco-2/HT29-MTX-E12 co-culture model system, we used the more *in vivo*-like human primary intestinal cells, the EpiIntestinal™ FT tissues. Due to the high costs of these tissues, only three mycotoxins, AOH, AME, and TeA at two concentrations (low and high) were chosen, and one experiment with two independent replicates was performed. Furthermore, the number of mucus-producing cells in the EpiIntestinal tissue is rather limited (Personal communication, 2024: Markus J, MatTek Life Sciences), and since the toxins’ effect on MUC5AC secretion in the co-culture was minor, mucus secretion was not measured in the EpiIntestinal tissues. Based on the co-culture experiments, the toxins alone had only minor effects on the cells; therefore, we focused on investigating whether the toxins exerted any modulatory effects on IL-1β–induced inflammation. As in the co-culture model, IL-1β (0.5 ng/mL) alone did not affect viability or the barrier of the EpiIntestinal™ FT tissue. Also, IL-1β, in combination with AOH (0.3 µM), and TeA (2, 100 µM), resulted in a small reduction in viability (metabolic activity) of about 6%, without affecting TEER values (**Fig. 7**).

**Fig 7:**
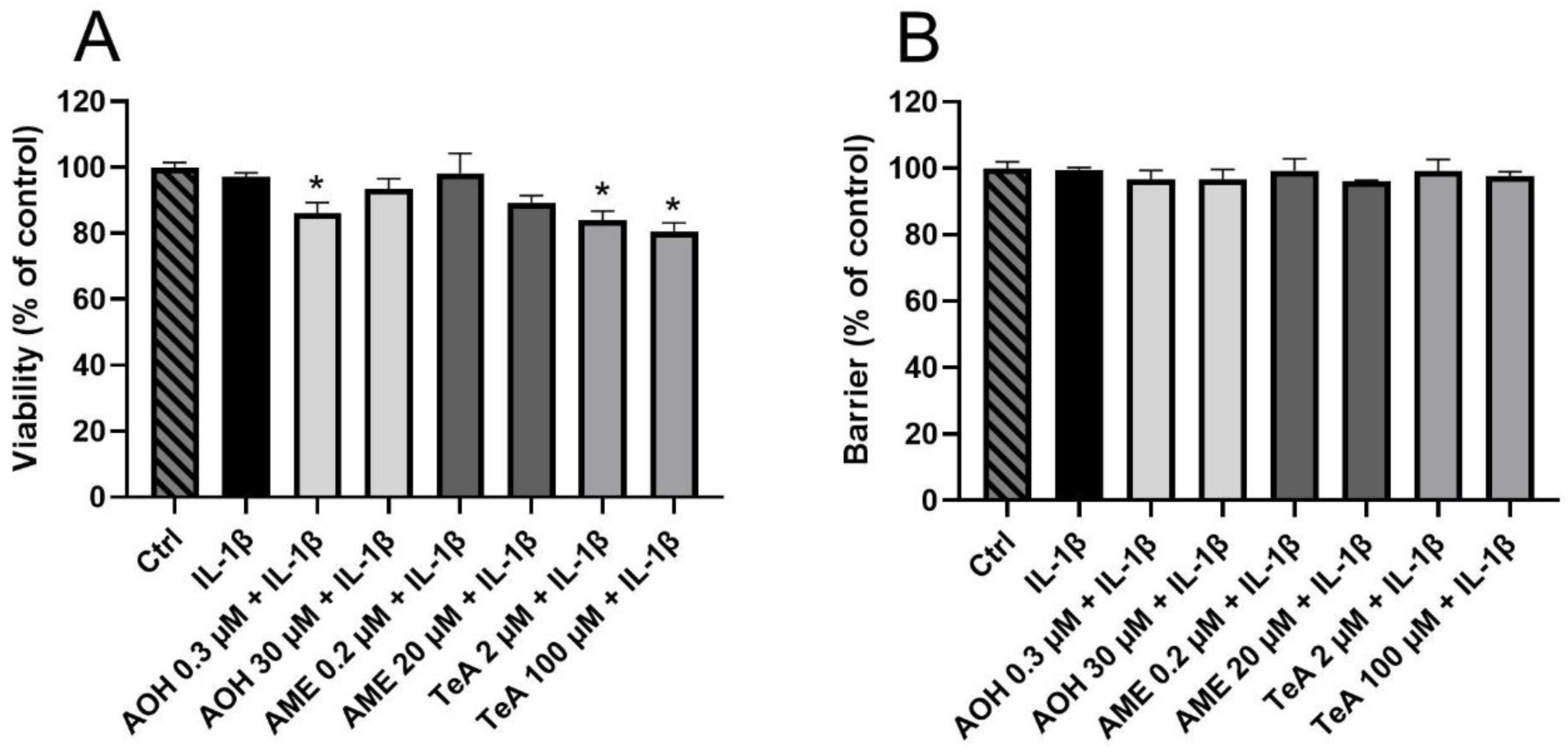
The EpiIntestinal™ FT tissues were treated with AOH, AME, and TeA for 2 h before the addition of IL-1β (0.5 ng/mL) for an additional 18 h, and viability / metabolic activity (**A**) and barrier integrity (**B**) were measured. The data are expressed as mean + SD from two independent incubations. Significant difference (p< 0.05) compared to IL-1β is indicated with an asterisk (*). One-way ANOVA with Dunnett’s post hoc test.

Next, we screened for effects of IL-1β (0.5 ng/mL) in combination with AOH (30 µM), AME (20 µM), or TeA (100 µM) on proinflammatory cytokines by simultaneously measuring secretion of IL-8, IL-6, TNFα, IL-1β, IL-10, and IL-12p70 using a cytometric bead array. Of these, only IL-8 and IL-6 showed elevated levels (**Supplementary, Fig S4**). Cytokine measurements were therefore repeated by using standard ELISA for IL-8 and IL-6 (**Fig. 8**). Here, IL-8 was induced by IL-1β, and in combination with TeA (100 µM), the induction was further increased. No significant effect was observed when IL-1β was combined with AOH or AME, but a tendency toward increased cytokine secretion was noted. While the induction of IL-8 by IL-1β was about 4-fold, the induction of IL-6 was about 10-fold. Although experimental variability was somewhat high, a significant increase in IL-6 was observed with AOH (30 µM) in combination with IL-1β.

**Fig. 8:**
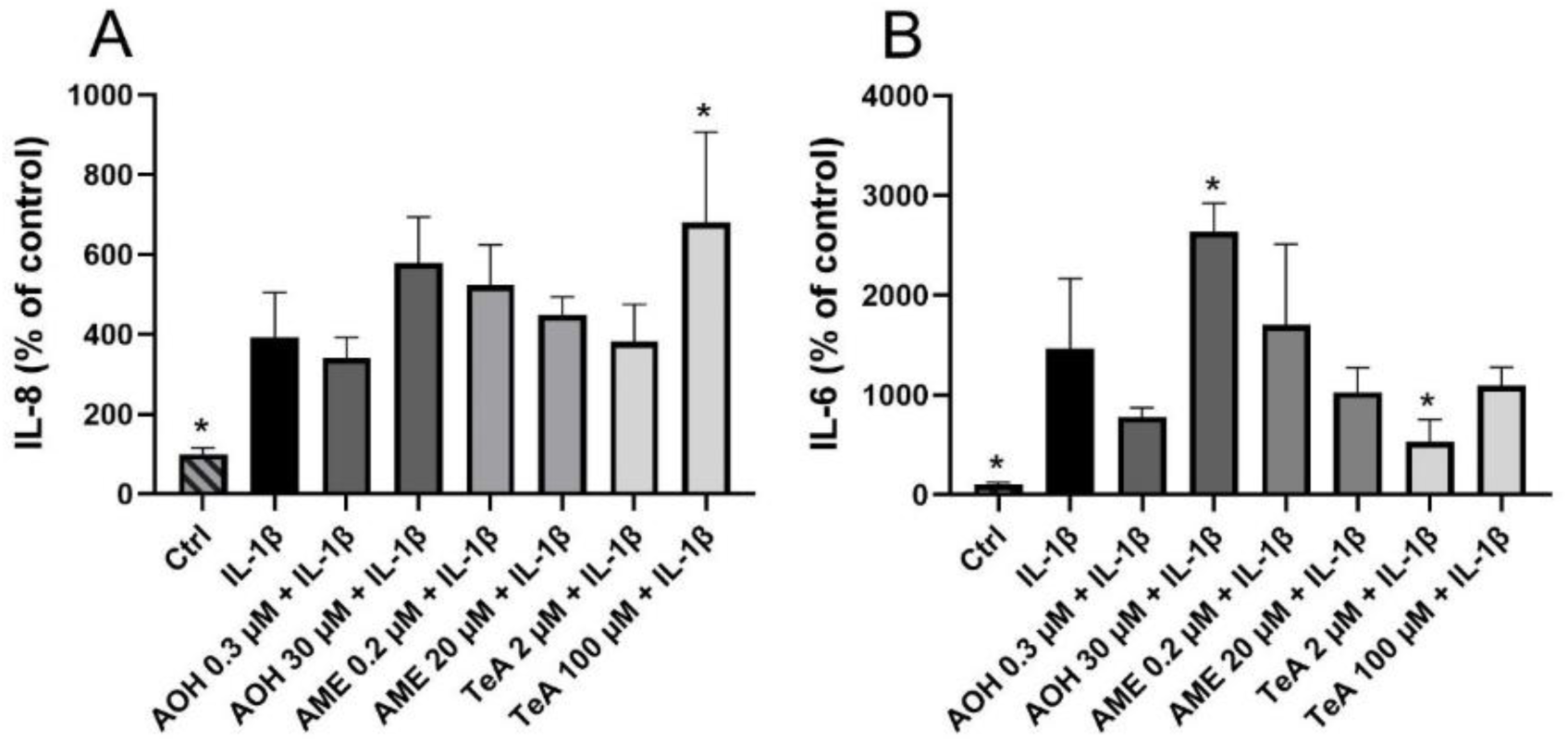
The EpiIntestinal*™* FT tissues were treated with AOH, AME, and TeA for 2 h before the addition of IL-1β (0.5 ng/mL) for an additional 18 h. IL-8 (**A**) and IL-6 (**B**) cytokine secretion was measured by ELISA. The data are expressed as mean + SD from two independent incubations, measured in triplicate. Significant difference (p< 0.05) compared to IL-1β is indicated with an asterisk (*). One-way ANOVA with Dunnett’s post hoc test.

### 3.3 Uptake studies of AOH, AME and TeA

The Caco-2 cell line has been widely used as a suitable *in vitro* model for predicting intestinal permeability *in vivo*. Generally, transport studies are recommended to be performed under linear conditions and at non-toxic concentrations. We therefore tested AOH, AME, and TeA at 10 µM rather than at the considerably higher concentrations at which biological effects were observed. Only the parental compounds were analysed for. Optimalisation experiments for uptake were performed using differentiated Caco-2 cells on transwell inserts and AOH (5 and 10 µM, for 0, 2, 4, 6, and 24 h). AOH was detected both intracellularly and in the basolateral compartment, indicating that AOH is taken up by the cells and released at the basolateral side (**Supplementary Fig. S6**), which represents the bloodstream. The AOH in the apical compartment decreases over time, while AOH in the basolateral compartment increases until 6 h and then levels out. Intracellular AOH first increases but then decreases after 2 h. Compared with the start of exposure to 10 µM AOH (0 h), the total remaining AOH (the sum across all compartments) decreased by 16% after 2 h, 43% after 6 h, and 90% after 24 h. The findings from the preliminary studies indicate that the model is useful and can be used with the more expensive, complex and more *in vivo*-like EpiIntestinal™ model. After 5 h of exposure to the EpiIntestinal™ tissue (**Fig. 9**), the uptake, the amount of toxin in the basolateral compartment (as % of total toxin distribution at the time of exposure), was highest for TeA (28 ± 2%), followed by AME (19 ± 9%) and AOH (8 ± 3%), while the intracellular concentration was highest for AME, followed by TeA and AOH, but still constituted a very low proportion of the total toxin distributions. The total remaining AOH, AME, and TeA (the sum of all compartments) are 22%, 13% and 74% of the measured starting concentrations, respectively, indicating metabolism and distinct toxicokinetic interactions or alternatively binding to plastics. A preliminary uptake experiment was also performed using differentiated Caco-2 cells, with higher total recoveries. Additionally, while AOH and AME levels in the basolateral compartment were about similar in both cell models, TeA was present at considerably higher levels in the basolateral compartment of EpiIntestinal tissue than in Caco-2 cells (**Supplementary Fig. S7**). Altogether, this indicates a difference between EpiIntestinal™ tissue and Caco-2 cells.

**Fig. 9:**
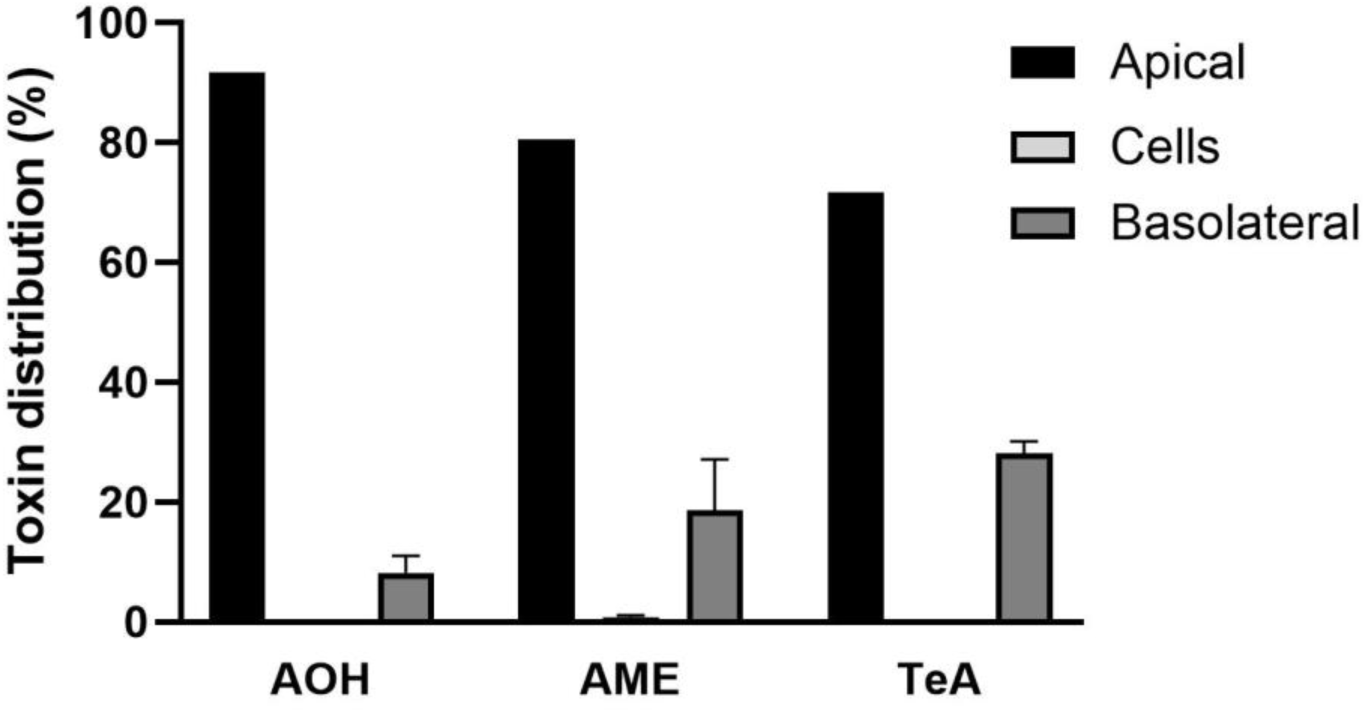
EpiIntestinalTM tissues were apically exposed to AOH (10 µM), AME (10 µM), or TeA (10 µM), followed by detection of the mycotoxins after 5 h, from the apical-, basolateral compartments, and in the cells. The data are expressed as mean + SD of three independent incubations.

Unwanted materials, such as mycotoxins, may be expelled by active transporters that are highly expressed in barrier cell types, including intestinal epithelial cells. Previously, it has been reported that several influx and efflux drug transporters are highly expressed in Caco-2 cells (Fedi et al. 2021; Maubon et al. 2007; Slattery et al. 2023). As a preliminary study, we used undifferentiated Caco-2 cells to investigate whether Alternaria mycotoxins interfere with the efflux transporter P-glycoprotein (P-gp), using the Calcein-AM assay. As a positive control, P-gp activity was inhibited with PSC833 (Kusunoki et al. 1998), which reduced P-gp-mediated efflux of calcein-AM, and thus increased the intracellular fluorescent calcein by about 100 % (**Supplementary Fig S8**). Only high concentrations of AME (40 µM) and ATX-1 (40 µM) had a minor, but not significant, effect, indicating that P-gp does not play a major role in regulation of uptake of Alternaria toxins. Nonetheless, follow-up studies using EpiIntestinal tissues, which express more intestinal-like features, with more normal expression of P-gp and other efflux pumps should be used to verify these findings.

## 4. Discussion

In this study, we have studied six of the most common Alternaria mycotoxins: AOH, AME, ALT, ATX-I, TEN, and TeA. using different *in vitro* intestinal cell models to examine barrier and immune responses of the Alternaria mycotoxins. Overall, we found that the toxins have only modest effects on human intestinal cells. Cytotoxicity was very low for all tested toxins, and consistent with this, none of them affected the intestinal barrier. While AOH, AME, and ATX-I induced cell cycle arrest and increased ROS levels, TeA reduced cell viability without causing cell cycle arrest or altering ROS levels. Furthermore, only AOH, AME, and TeA slightly modulated proinflammatory cytokine secretion in combination with IL-1β.

Although a wide range of concentrations is used in this study, the amount of mycotoxins that may reach the intestine after consuming food contaminated with Alternaria mycotoxins varies widely. The highest exposure to AOH for toddlers is estimated to be 3.8 - 38.8 ng/kg bw/day, with a 95^th^ percentile of 11.4 - 270.5 ng/kg bw/day. The exposure to AME is almost equal to AOH, while exposure to TeA ranges from 100 to 1614 ng/kg bw/day, with a 95^th^ percentile of 98 - 3603 ng/kg bw/day (EFSA 2016). Based on this, exposure to these toxins through three daily meals for a toddler with a weight of 20 kg, one meal can be estimated to contain up to 1.8 µg AOH and AME, and 10.8 µg TeA. Assuming a stomach volume of 0.5 L for a toddler, this would correspond to approximately 0.014 µM AOH and AME, and 0.05 µM TeA, which is concentrated about 10-fold during the reabsorption of 90% of the water as it passes through the upper parts of the gastrointestinal tract. The lower concentrations used in our experiments (AOH 0.3-60 µM, AME 0.2–40 µM, and TeA 2–200 µM) are therefore within a realistic range for the lower parts of the intestine. The estimated intake of TEN is 1.6–33.4 ng/kg bw/day (EFSA 2016) and is thus even lower than that of AOH. For the other Alternaria toxins used in this study, ALT and ATX-I, both intake and occurrence data are limited. It is difficult to estimate real-life exposure because exposure levels vary and occurrence data for several of the Alternaria mycotoxins are scarce. Also, several mycotoxins co-exist, and they may influence each other’s effects (Louro et al. 2024; Solhaug et al. 2016b). Nonetheless, for modelling potential hazards in an *in vitro* system, the concentrations used in this study are judged by us to be within the relevant concentration range. The lower concentrations may represent a realistic exposure range, and the higher concentrations are more to explore the potential toxicity.

Proliferating Caco-2 cells are often considered a model of intestinal crypt cells, whereas fully differentiated Caco-2 monolayers closely resemble villus enterocytes in both morphology and function, representing mature enterocyte-like cells. Compared with differentiated cultures, proliferating Caco-2 cells appear more susceptible to external stressors (Bu et al. 2016). This increased vulnerability may be partly explained by their high rate of cell division. Because these cells undergo continuous DNA replication, they are more prone to replication stress and the accumulation of DNA damage, which can lead to cell cycle arrest, apoptosis, or reduced cell viability following exposure to genotoxic agents. Cell cycle arrest is often linked to DNA damage. Numerous studies, including both immune cells (Solhaug et al. 2013; Solhaug et al. 2016b) and intestinal cells (Fernandez-Blanco et al. 2016; Marin et al. 2024), have shown that AOH and AME induce cell cycle arrest, most often in the G2 phase. The mechanism is coupled to their ability to act on topoisomerase II (Fehr et al. 2009), and the following DNA damage response pathways, involving γH2AX, p53 and Cyclin B (Solhaug et al. 2016a). Less is known about the DNA damage potential of the perylene quinone ATX-I. But it has tested positive for double-stranded DNA breaks using the alkaline unwinding assay (Louro et al. 2024) and shown to interfere with topoisomerase II (Jarolim et al. 2017). While the genotoxic potential of ATX-I remains to be elucidated further, it is found here, along with AOH and AME, at a similar concentration range, to reduce Caco-2 cell proliferation, with a significant decrease of cells in the G1 phase and an apparent accumulation of cells in both S- and G2 phases. If the cell cycle arrest in dividing Caco-2 cells reflects what happens in dividing enterocytes in the crypts *in vivo,* this indicates a potential toxic effect on the gastrointestinal epithelium. In contrast to Caco-2 cells, AOH induces a clearer G2 arrest in THP-1 monocytes (Solhaug et al. 2016b) and RAW264.7 cells (Solhaug et al. 2012). One explanation can be that the Caco-2 cell line has several mutations in the important cell cycle regulator p53 (Bairoch 2018). Despite oncogenic pathway alterations, Caco-2 cells undergo spontaneous G0/G1 arrest upon differentiation in long-term culture (Ding et al. 1998), illustrating strong context-dependence of proliferation control. Nevertheless, cell cycle progression is clearly delayed, indicating DNA damage. In the absence of functional p53, cells often avoid apoptosis after DNA damage because they cannot activate the intrinsic mitochondrial death pathway, which may explain why the Caco-2 cells do not undergo cell death even after 72 h of exposure. This may also be an explanation for the ability of AOH to induce apoptosis in the p53wt human colorectal carcinoma cell line, HCT116 (Bensassi et al. 2012). Interestingly, TeA clearly inhibited the metabolic activity of Caco-2 cells but had no effect on the cell cycle. TeA inhibits protein synthesis by blocking the peptidyl transferase reaction on the 60S ribosomal subunit (Vazquez et al. 1974). This reduces the protein production, which is essential for proper regulation and progression of cell proliferation. Since the cells did not accumulate in any specific cell cycle phase, this suggests a general slowdown in cell cycle progression. So far, there are no reports of TeA being genotoxic (Louro et al. 2024), which supports the lack of a specific cell-cycle arrest observed in Caco-2 cells. Very few studies have examined the effects of ALT (Louro et al. 2024). A study using the HCT116 cell line found that ALT induced cytotoxicity after 72 h, with an IC_50_ of 3.13 µM (Xiao et al. 2014), whereas no cytotoxicity was detected in the keratinocyte cell line HaCaT after 24 h of exposure at concentrations up to 80 µM (Dong et al. 2021). The mechanism of action of ALT is still unknown, but it is negative in bacterial mutagenicity assays (Schrader et al. 2006; Schrader et al. 2001), which supports our data showing no effects on the cell cycle and only a minor decrease of 12% in metabolic activity after 72 h at the highest concentration tested (40 µM). Furthermore, TEN did not affect neither metabolic activity nor the cell cycle, consistent with experiments on the HepaGR cell line (Hessel-Pras et al. 2019), in which no cytotoxicity was detected.

Oxidative stress results from an imbalance between oxidants and antioxidants and is primarily driven by excessive ROS production and impaired oxidative detoxification pathways. Here, AOH, AME, and ATX-I also caused cell cycle arrest and increased the cellular ROS levels. Consistent with previous studies, both AOH and AME are found to enhance the ROS level and to affect ROS-associated cell signalling pathways (Fernandez-Blanco et al. 2014; Marin et al. 2024; Solhaug et al. 2012; Tiessen et al. 2013). However, the exact mechanism of how AOH and AME produce ROS is not known. Both AOH and AME have been shown to undergo aromatic hydroxylation (Borsos et al. 2024; Pfeiffer et al. 2007), thereby generating reactive catechols and hydroxyquinones that may generate ROS during quinone redox cycling. Also, because ATX-I is a perylene quinone, semiquinone formation is chemically possible, but has not, to the authors’ knowledge, yet been demonstrated. However, ROS is primarily produced in mitochondria due to incomplete reduction of oxygen during oxidative phosphorylation for ATP production. Defects in mitochondrial function, such as inefficient oxidative phosphorylation and loss of membrane potential, can lead to elevated ROS production. It is therefore also possible that the elevated ROS level arises from cellular stress as a part of the DNA-damaging response.

The intestine is the first physiological barrier against food contaminants. Several mycotoxins, such as aflatoxins, ochratoxins, and deoxynivalenol, are known to impair intestinal permeability (Gao et al. 2020). Factors that can affect the barrier include acute cytotoxicity, reduced expression of tight junction proteins, and inflammation. A defective intestinal tight junction barrier is an important contributor to the pathogenesis of various inflammatory gut conditions, including Inflammatory bowel diseases (IBDs) (Kaminsky et al. 2021). None of the Alternaria toxins tested here affected the barrier, neither alone nor in combination with the proinflammatory cytokine IL-1β. Interestingly, only the highest TeA concentration (100 µM) reduced metabolic activity in the Caco-2/HT29-MTX-E12 co-culture, but without affecting the barrier. This suggests that TeA only has a minor effect on cell viability. In contrast, this reduction was not observed when IL-1β was combined with TeA in the co-culture. Consistent with this, TeA in combination with IL-1β only reduced the viability by about 6% in the EpiIntestinal™ tissue. The ability of IL-1β to counteract this effect may indicate that it activates inflammatory signalling that upregulates metabolic pathways or stress responses that compensate for the effect of TeA.

Proinflammatory cytokines, including IL-1β, are markedly elevated in the gut during inflammatory conditions. IL-1β is mainly secreted by activated macrophages, and its actions are primarily mediated by NF-κB activation, leading to increased intestinal barrier permeability and the secretion of proinflammatory cytokines (Kaminsky et al. 2021). While intestinal epithelial cells are not considered part of the immune system, they can produce several cytokines and chemokines that are crucial for recruiting and activating immune cells, such as lymphocytes and macrophages. Enterocytes secrete the chemokine IL-8 to attract and activate neutrophils during the acute phase of inflammation, while IL-6 is a cytokine that mediates pleiotropic effects during both acute and chronic inflammation. In Caco-2 cells, IL-1β is found to be more potent than TNFα, IFNγ, and LPS in inducing IL-8 secretion and IL-6 production (Van De Walle et al. 2010). So far, most work on Alternaria toxins has focused mainly on immune cells, and studies indicate that some of them, AOH, AME, ATX-I, ALT and TEN, but not TeA, can act as immunosuppressors. The immunosuppressive effect is shown to involve interference with the transcription factor NF-κB and thus leads to a decrease in the secretion of proinflammatory cytokines (Crudo et al. 2025; Dong et al. 2021; Grover and Lawrence 2017; Kollarova et al. 2018; Liu et al. 2024; Louro et al. 2024). So far, few studies have been done on intestinal cells, but Schmutz and co-workers reported minor changes in IL-8 and IL-1 gene expression in response to AOH in Caco-2 cells (Schmutz et al. 2019). In agreement with this, none of the Alternaria toxins tested in this study affected IL-8 secretion in unstimulated Caco-2 cells. In contrast, AOH can induce secretion of both IL-6 and TNF-α, but not IL-8, IL-10, and IL-12p70 in human macrophages (Solhaug et al. 2015). In our study using the Caco-2/HT29-MTX-E12 co-culture, the highest concentration of AOH (30 µM) and AME (20 µM) reduced IL-1β-induced (0.5 ng/mL) IL-8 secretion, whereas the lowest concentration of AME (0.2 µM) surprisingly enhanced the IL-8 secretion to some extent. Also, in Schmutz and co-workers, higher concentrations of AOH (20 and 40 µM) slightly reduced IL-1β (25 ng/mL)-induced IL-8 secretion, whereas a lower concentration of AOH (0.2 µM) enhanced it (Schmutz et al. 2019). Furthermore, (Partsch et al. 2026) observed a downregulation of IL-1β (25 ng/mL) induced IL-8 upon exposure of Caco-2 to AOH (30 µM) and AME (10 µM). Interestingly, in the EpiIntestinal™ tissue, which can be considered a more realistic *in vivo*-like model than the Caco-2/HT29-MTX-E12 co-culture, these inhibitory effects on IL-1β-induced IL-8 secretion were not observed. Here, it seems more of a tendency that AOH (30 µM) enhanced IL-8 and IL-6 secretion induced by IL-1β (0.5 ng/mL). Interestingly, the highest concentration of TeA (100 µM) also enhanced IL-1β-induced IL-8 secretion in both the co-culture and the EpiIntestinal™ tissue but had no effect on the IL-6 secretion. A comparison of the results from EpiIntestinal™ tissue with the co-culture model indicates that the co-culture model underestimates the proinflammatory responses of AOH in combination with IL-1β. It is difficult to explain these distinctive results, but a plausible explanation is that the toxins may act on different signalling pathways in the different cell models. While both AOH and AME are believed to affect the NF-κB signalling pathway, TeA had no effect on this pathway (Crudo et al. 2025; Partsch et al. 2026). Interestingly, the ERK1/2 pathway is shown to mediate IL-8 secretion independently of NF-κB activation in response to cadmium in lung epithelial cells (Cormet-Boyaka et al. 2011). It would therefore be interesting to elucidate the effects of the Alternaria toxins, especially TeA, on this pathway as well. The other toxins tested, ATX-I, TEN, and ALT, had no effect on IL-8 secretion in the co-culture and were therefore not included in the experiments using the EpiIntestinal™ tissue. So far, only a very limited number of immunological studies have been performed on these compounds. In a study using skin keratinocytes (HaCaT), ALT had no effect on INFγ/TNFα-induced IL-6 secretion (Dong et al. 2021). Furthermore, in THP-1 monocytes and macrophages, AME, ATX-I, TEN and ALT were recently found to reduce NF-kB activity. AME, however, only at cytotoxic concentrations (Partsch et al. 2026). Overall, these compounds appear to have a very low immunotoxic potential. Cultivating cells with flowing cultivating medium simulating the circulating bloodstream may affect the cells and their response to external stimuli. It is known that cultivation of Caco-2 cells with a flow may affect the polarization and morphology of the monolayer (Delon et al. 2019; Fois et al. 2021) and, therefore, also may affect the immune response. However, in our co-culture model, only minor differences between results with and without flow were observed, and we concluded that flow added limited value to this study.

The mucus lining the intestinal epithelium provides an important physicochemical barrier protecting against ingested pathogens and toxins (Grondin et al. 2020). Goblet cells in the intestinal epithelium synthesize and secrete gel-forming mucins, including MUC2 and MUC5AC, which contribute to the mucus layer (Pelaseyed et al. 2014). Although MUC2 is the dominant secreted mucin in the intestine, MUC5AC can be induced under inflammatory or injury-related conditions (Olli et al. 2020). The rapid elevation of secretory mucin in response to xenobiotics and bacterial infection is crucial for protecting the intestine. Interestingly, the Fusarium mycotoxin deoxynivalenol was found to upregulate mRNA expression of MUC5AC, MUC5B, MUC4 and MUC17 in Caco-2 (Wan et al. 2025), and increase the MUC5AC and MUC5B secretion in the Caco-2/HT29-MTX co-culture (Wan et al. 2014). While several studies have shown that mycotoxins can affect the mucus layer (Robert et al. 2017; Wan et al. 2014), this is the first study that includes Alternaria mycotoxins. Even though expression of several mucins, including MUC5AC, is suggested to be activated by the NF-κB signalling pathway (Li et al. 2021), AOH, AME, TEN and ATX-I, which are reported to suppress this pathway (Partsch et al. 2026), had no effect on MUC5AC secretion in the Caco-2 /HT29-MTX-E12 co-culture. Interestingly, in our study, a low concentration (0.2 µM) of ALT increased MUC5AC secretion by 45% compared to control. This may suggest that other signalling pathways besides NF-κB are involved, such as the PKCδ/θ - Ras-Raf-MEK/ERK signalling pathway, which is identified to be important for MUC5AC induction in lung epithelial cells (Hewson et al. 2004).

Absorption is essential for the bioavailability of ingested compounds. It is partly dependent on the compounds’ lipophilicity, or affinity for lipid environments. Higher lipophilicity enhances passive diffusion by increasing their segregation into the phospholipid bilayer. However, excessive lipophilicity can also reduce solubility. Less lipophilic compounds can be transported over the epithelial barrier paracellularly, between the tight junctions (Chmiel et al. 2019). The uptake, transport from the apical to the basolateral compartment of AOH, AME, and TeA (all 10 µM) in EpiIntestinal™ tissue was quite low, with the highest amount was for TeA (28 ± 2%), followed by AME (19 ± 9%) and AOH (8 ± 3%). However, the total recovery of the paternal compounds, AOH, AME and TeA in the sum of the three analysed compartments, apical, basolateral and in the cells, were 22, 13 and 74% respectively. The difference in the preliminary estimates of the apparent permeability coefficient (P_app_) values in the EpiIntestinal^TM^ model based on disappearance from the apical side vs increase on the basolateral or receiving side (7,4 × 10^-5^ vs 3,1 × 10^-7^ cm/s for AOH and 8,3 × 10^-5^ vs 4,4 × 10^-7^ cm/s for AME and 4,3 × 10^-5^ vs 3,9 × 10^-6^ cm/s for TeA, ) together with the low total recovery indicates that the toxins are extensively metabolised in the gastrointestinal tissue prior to absorption as this model is documented to have a considerably higher biotransformation capacity than the Caco-2 cell model (Cui et al. 2020). The EpiIntestinal model has been shown to have a high potential to predict human absorption (Chothe et al. 2025), and P_app_ values above 5 × 10^-6^ in the EpiIntestinal^TM^ model were classified as high permeability substances (Markus et al. 2021). Our findings are an indication of absorption from the gastrointestinal tract, a finding that fits well with the reported findings of all these Alternaria toxins in urine (Qiao et al. 2020). In studies of occurrence in human urine, however, the exposure to the Alternaria toxins is uncertain, and it is therefore difficult to assess the uptake based on those studies. Furthermore, the urine samples were analysed for conjugated and unconjugated toxins by use of deconjugation enzymes. This approach does not take potential other metabolites into account. Data from the EpiIntestinal^TM^ on absorption of drugs have been demonstrated to be well adapted to PBPK modelling of drugs (Chothe et al. 2025), and could be used in future PBPK modelling these toxins. An alternative explanation for the low total recovery could be binding to the plastics, but since this was not observed in studies of the considerably more lipophilic enniatins; LogP 4.1 – 4.7 for enniatin A, A1 and B1 and 3.9 for enniatin B1, vs 3.8, 3.9 and 0.57 for AOH, AME and TeA, (Sdogati et al. 2026; Tran et al. 2020) and as personal experience (Solhaug et al. 2026, unpublished results), we consider this to be less likely.

A study investigated the metabolism and excretion of 17 Alternaria toxins in rats (Puntscher et al. 2019). Out of the administered doses of AOH, AME, and TeA (up to 12, 10, and 8.945 μg/kg bw, respectively), 6 – 10% / 9%, - 6-7% / 87%, and 55% / 0.3% were recovered in urine and faeces during 24 h, respectively. Both AOH and AME can undergo both oxidoreductive phase I metabolism and conjugative phase II metabolism (Louro et al. 2024). In agreement with this, AOH is likely to be extensively metabolised as the total recovery of AOH in urine and faeces was below 20% (Puntscher et al. 2019). Furthermore, *in vitro* studies using Caco-2 cells, Burkhardt et al. (2009) found that about 25% of AOH added apically reached the basolateral compartment after 3 h of exposure, either as AOH or as its metabolites: AOH-3-O-glucuronide, AOH-9-O-Glucuronide, and AOH-3-O-Sulfate. Metabolism may also explain why we found only 8 ± 3% of AOH in the basolateral compartment. In contrast to AOH, 87% of the AME was found unchanged in the faeces; which indicates that absorption of AME and/or its metabolism is probably very low in rats (Puntscher et al. 2019). Burkhardt et al., (2009) also reported low AME absorption in Caco-2 cells, with about 3-7% of the added toxin reaching the basolateral compartment, mainly as AME-3-O-glucuronide. In EpiIntestinalTM tissue, we found somewhat higher AME levels (19 ± 9%) in the basolateral compartment, indicating higher absorption than in other studies. Also, the metabolites were not analysed here, which may have led to even higher levels, but no conjugates of AME were detected in human urine samples (Qiao et al., 2020), indicating a limited significance of conjugated metabolites. An alternative explanation for the difference in these results may be the different experimental models, as we have used the EpiIntestinal™ tissue consisting of human primary cells and not a carcinoma-derived cell line. Animal and human studies indicate that TeA is excreted with urine as the major excretion route (Louro et al. 2024; Puntscher et al. 2019). So far, no *in vitro* data on the metabolism of TeA are available, but data from a human *in vivo* study indicate that TeA is apparently not metabolised or has a high metabolic stability (Louro et al. 2024). Interestingly, we found that after 5 hr of exposure of the EpiIntestinal™ tissues, the total remaining amount of TeA was 74% of the measured starting concentration, and 28 ± 2% was found in the basolateral compartment and almost nothing in the cells, which does not exclude the possibility that TeA is metabolised in our system. Altogether, our findings indicate that the tested compounds differ in several toxicokinetic parameters, including metabolism. However, this could be investigated further by metabolite profiling.

In addition to lipophilicity, efflux pumps that actively transport certain compounds back into the intestinal lumen reducing intracellular concentrations, are important for absorption. While the Alternaria mycotoxins, AOH, AME, TeA, ATX-I, TEN, and ALT did not interact with the most prominent efflux transporter, P-gp, in our study, it does not exclude their interaction with other efflux pumps, such as Breast Cancer Resistance Protein (BCRP) and Multidrug Resistance Protein 2 (MRP2). However, this has not yet been studied. Interestingly, a recent study has identified AOH as both a substrate and an inhibitor of influx transporters, the Solute Carrier Organic Anion Transporting Polypeptides (OATPs) (Fliszár-Nyúl et al. 2023). It is well established that certain OATPs can influence cellular uptake. AOHs interaction with OATPs therefore suggests that uptake of AOH is influenced by OATPs and that this may also affect the uptake of other endogenous and exogenous compounds, including other mycotoxins and drugs where the uptake is influenced by these transporters during co-exposure. Furthermore, the observed differences in the accumulation of the tested mycotoxins in cells may partly be explained by differences in their affinity for OATPs. Our preliminary uptake study using Caco-2 cells, presented in the supplementary material, shows some differences compared to the EpiIntestinal™ tissue. Important factors that may have influenced this difference include the Caco-2 model’s overexpression of tight junctions, which makes paracellular transport of less lipophilic compounds less comparable. Furthermore, there are considerable differences in the expression of transporters such as P-gp, and metabolising enzymes responsible for biotransformation (e.g. CYP3A4). Also, the absence of a mucus layer in the Caco-2 model is considered a major drawback as it can facilitate the absorption of less lipophilic compounds (Cui et al. 2020; Fedi et al. 2021; Markus et al. 2021). As a consequence of this, EpiIntestinal™ tissue discriminated between low- and high-permeability drugs and correlated more strongly with human absorption data (r2 = 0.91) than Caco-2 cells (r^2^ = 0.71) (Ayehunie et al. 2018).

The existing knowledge on toxicity of Alternaria toxins has been reviewed and lack of data needed to complete a risk assessment have been identified (EFSA 2011; Louro et al. 2024). Among other data gaps, it was pointed out that there is a lack of information on ADME properties as well as on immunotoxicological effects other than cell lines. In addition, the reviews did not identify any information on the potential effects on the gastrointestinal tract for the Alternaria toxins with the exception of some *in vitro* findings on potential inflammatory effects on the gastrointestinal Caco-2 cell line. Our findings from a human relevant model indicate that the effect on intestinal barrier function is limited. Similarly, Alternaria toxins only triggered a proinflammatory response in the human relevant GI model at rather high concentrations compared to human dietary exposure. In conclusion, the studied Alternaria toxins have a moderate effect on the intestinal tract *in vitro*, but only at concentrations exceeding those expected from normal food intake. We observed mild inflammatory effects in the gastrointestinal models used here. The immunomodulatory effects also differed between the Caco-2 cells, the Caco-2/HT29-MTX-E12 co-culture model, and the human primary cell model, underlining the need for a relevant model for the study’s purpose and endpoints. We also conclude that the gastrointestinal absorption of AOH is limited, while the absorption of TeA is higher. For AME and TeA, further studies of the absorption are needed, including analysis of potential metabolites and studies of human toxicokinetic parameters like maximum plasma concentrations and half-life and excretion are warranted.

## Supporting information

Supplementary_Solhaug et al

## Acknowledgement

This work was carried out in the framework of the European Partnership for the Assessment of Risks from Chemicals (PARC) and has received funding from the European Union’s Horizon Europe research and innovation programme under Grant Agreement No 101057014. Views and opinions expressed are however those of the author(s) only and do not necessarily reflect those of the European Union or the Health and Digital Executive Agency. Neither the European Union nor the granting authority can be held responsible for them. The work was co-funded by the Norwegian Veterinary Institute. Vanessa Partsch was a visiting PhD student from the University of Vienna, and her stay was funded by the Vienna Doctoral School in Chemistry (DoSChem). The graphical abstract was created in BioRender. Solhaug, A. (2026) https://BioRender.com/33kmgwv. We also wish to thank Cristin Plassen, Norwegian Veterinary Institute, for her skilful help in the lab.

## References

Aichinger G, Del Favero G, Warth B, Marko D (2021) Alternaria toxins—Still emerging? Comprehensive Reviews in Food Science and Food Safety 20(5):4390–4406 10.1111/1541-4337.12803

Al-Sadi RM, Ma TY (2007) IL-1beta causes an increase in intestinal epithelial tight junction permeability. Journal of immunology 178(7):4641–9 10.4049/jimmunol.178.7.4641

Ayehunie S, Landry T, Stevens Z, Armento A, Hayden P, Klausner M (2018) Human Primary Cell-Based Organotypic Microtissues for Modeling Small Intestinal Drug Absorption. Pharmaceutical research 35(4):72 10.1007/s11095-018-2362-0

Bairoch A (2018) The Cellosaurus, a Cell-Line Knowledge Resource. J Biomol Tech 29(2):25–38 10.7171/jbt.18-2902-002

Bensassi F, Gallerne C, El Dein OS, Hajlaoui MR, Bacha H, Lemaire C (2012) Cell death induced by the Alternaria mycotoxin Alternariol. Toxicology in Vitro 26(6):915–923 10.1016/j.tiv.2012.04.014

Borsos E, Varga E, Aichinger G, Marko D (2024) Unraveling Interspecies Differences in the Phase I Hepatic Metabolism of Alternariol and Alternariol Monomethyl Ether: Closing Data Gaps for a Comprehensive Risk Assessment. Chemical research in toxicology 37(8):1356–1363 10.1021/acs.chemrestox.4c00095

Bu P, Narayanan S, Dalrymple D, Cheng X, Serajuddin AT (2016) Cytotoxicity assessment of lipid-based self-emulsifying drug delivery system with Caco-2 cell model: Cremophor EL as the surfactant. Eur J Pharm Sci 91:162–71 10.1016/j.ejps.2016.06.011

Burkhardt B, Pfeiffer E, Metzler M (2009) Absorption and metabolism of the mycotoxins alternariol and alternariol-9-methyl ether in Caco-2 cells in vitro. Mycotoxin Res 25(3):149–57 10.1007/s12550-009-0022-2

Chmiel T, Mieszkowska A, Kempińska-Kupczyk D, Kot-Wasik A, Namieśnik J, Mazerska Z (2019) The impact of lipophilicity on environmental processes, drug delivery and bioavailability of food components. Microchemical Journal 146:393–406 10.1016/j.microc.2019.01.030

Chothe PP, Whitcher-Johnstone A, Karve A, Ramsden D, Hariparsad N (2025) Advancing Predictions of Oral Drug Absorption, CYP3A4 Induction, and Transporter-Mediated Interactions Using a Human Primary Intestinal 3D Model (EpiIntestinal™). Clin Pharmacol Ther 118(6):1499–1511 10.1002/cpt.70000

Cormet-Boyaka E, Jolivette K, Bonnegarde-Bernard A, et al. (2011) An NF-κB–Independent and Erk1/2-Dependent Mechanism Controls CXCL8/IL-8 Responses of Airway Epithelial Cells to Cadmium. Toxicological Sciences 125(2):418–429 10.1093/toxsci/kfr310

Crudo F, Partsch V, Braga D, et al. (2025) Discovery of the Alternaria mycotoxins alterperylenol and altertoxin I as novel immunosuppressive and antiestrogenic compounds in vitro. Archives of toxicology 99(1):407– 421 10.1007/s00204-024-03877-1

Cui Y, Claus S, Schnell D, Runge F, MacLean C (2020) In-Depth Characterization of EpiIntestinal Microtissue as a Model for Intestinal Drug Absorption and Metabolism in Human. Pharmaceutics 12(5) 10.3390/pharmaceutics12050405

Del Favero G, Hohenbichler J, Mayer RM, Rychlik M, Marko D (2020) Mycotoxin Altertoxin II Induces Lipid Peroxidation Connecting Mitochondrial Stress Response to NF-κB Inhibition in THP-1 Macrophages. Chemical research in toxicology 33(2):492–504 10.1021/acs.chemrestox.9b00378

Delon LC, Guo Z, Oszmiana A, et al. (2019) A systematic investigation of the effect of the fluid shear stress on Caco-2 cells towards the optimization of epithelial organ-on-chip models. Biomaterials 225:119521 10.1016/j.biomaterials.2019.119521

Ding Q-M, Ko TC, Evers BM (1998) Caco-2 intestinal cell differentiation is associated with G1 arrest and suppression of CDK2 and CDK4. American Journal of Physiology-Cell Physiology 275(5):C1193– C1200 10.1152/ajpcell.1998.275.5.C1193

Dong L, Kim HJ, Cao TQ, et al. (2021) Anti-Inflammatory Effects of Metabolites from Antarctic Fungal Strain Pleosporales sp. SF-7343 in HaCaT Human Keratinocytes. International journal of molecular sciences 22(18):9674 10.3390/ijms22189674

EFSA (2011) Scientific Opinion on the risks for animal and public health related to the presence of Alternaria toxins in feed and food. EFSA Journal 9(10):2407 10.2903/j.efsa.2011.2407

EFSA (2016) Dietary exposure assessment to Alternaria toxins in the European population. EFSA Journal 14(12):e04654 10.2903/j.efsa.2016.4654

Fedi A, Vitale C, Ponschin G, Ayehunie S, Fato M, Scaglione S (2021) In vitro models replicating the human intestinal epithelium for absorption and metabolism studies: A systematic review. J Control Release 335:247–268 10.1016/j.jconrel.2021.05.028

Fehr M, Pahlke G, Fritz J, et al. (2009) Alternariol acts as a topoisomerase poison, preferentially affecting the IIα isoform. Mol Nutr Food Res 53(4):441–451 10.1002/mnfr.200700379

Fernandez-Blanco C, Font G, Ruiz MJ (2014) Oxidative stress of alternariol in Caco-2 cells. Toxicol Lett 229(3):458–64 10.1016/j.toxlet.2014.07.024

Fernandez-Blanco C, Juan-Garcia A, Juan C, Font G, Ruiz MJ (2016) Alternariol induce toxicity via cell death and mitochondrial damage on Caco-2 cells. Food and chemical toxicology : an international journal published for the British Industrial Biological Research Association 88:32–9 10.1016/j.fct.2015.11.022

Fleck SC, Burkhardt B, Pfeiffer E, Metzler M (2012) Alternaria toxins: Altertoxin II is a much stronger mutagen and DNA strand breaking mycotoxin than alternariol and its methyl ether in cultured mammalian cells. Toxicology Letters 214(1):27–32 10.1016/j.toxlet.2012.08.003

Fliszár-Nyúl E, Ungvári O, Dombi Á, Özvegy-Laczka C, Poór M (2023) Interactions of Mycotoxin Alternariol with Cytochrome P450 Enzymes and OATP Transporters. Metabolites 13(1):45 10.3390/metabo13010045

Fois CAM, Schindeler A, Valtchev P, Dehghani F (2021) Dynamic flow and shear stress as key parameters for intestinal cells morphology and polarization in an organ-on-a-chip model. Biomedical Microdevices 23(4):55 10.1007/s10544-021-00591-y

Gao Y, Meng L, Liu H, Wang J, Zheng N (2020) The Compromised Intestinal Barrier Induced by Mycotoxins. Toxins 12(10) 10.3390/toxins12100619

Gollapudi S, Kim CH, Tran B-N, Sangha S, Gupta S (1997) Probenecid reverses multidrug resistance in multidrug resistance-associated protein-overexpressing HL60/AR and H69/AR cells but not in P-glycoprotein-overexpressing HL60/Tax and P388/ADR cells. Cancer Chemotherapy and Pharmacology 40(2):150–158 10.1007/s002800050640

Grondin JA, Kwon YH, Far PM, Haq S, Khan WI (2020) Mucins in Intestinal Mucosal Defense and Inflammation: Learning From Clinical and Experimental Studies. Frontiers in Immunology Volume 11–2020 10.3389/fimmu.2020.02054

Grover S, Lawrence CB (2017) The Alternaria alternata Mycotoxin Alternariol Suppresses Lipopolysaccharide-Induced Inflammation. International journal of molecular sciences 18(7) 10.3390/ijms18071577

Hessel-Pras S, Kieshauer J, Roenn G, Luckert C, Braeuning A, Lampen A (2019) In vitro characterization of hepatic toxicity of Alternaria toxins. Mycotoxin Research 35(2):157–168 10.1007/s12550-018-0339-9

Hewson CA, Edbrooke MR, Johnston SL (2004) PMA induces the MUC5AC respiratory mucin in human bronchial epithelial cells, via PKC, EGF/TGF-alpha, Ras/Raf, MEK, ERK and Sp1-dependent mechanisms. Journal of molecular biology 344 3:683–95 10.1016/j.jmb.2004.09.059

Ivanova L, Uhlig S, Eriksen G, Johannessen L (2010) Enniatin B1 is a substrate of intestinal P-glycoprotein, multidrug resistance-associated protein 2 and breast cancer resistance protein. World Mycotoxin Journal 3(3):271–281 10.3920/WMJ2010.1225

Jarolim K, Del Favero G, Ellmer D, et al. (2017) Dual effectiveness of Alternaria but not Fusarium mycotoxins against human topoisomerase II and bacterial gyrase. Archives of toxicology 91(4):2007–2016 10.1007/s00204-016-1855-z

Kaminsky LW, Al-Sadi R, Ma TY (2021) IL-1β and the Intestinal Epithelial Tight Junction Barrier. Frontiers in Immunology Volume 12–2021 10.3389/fimmu.2021.767456

Kollarova J, Cenk E, Schmutz C, Marko D (2018) The mycotoxin alternariol suppresses lipopolysaccharide-induced inflammation in THP-1 derived macrophages targeting the NF-κB signalling pathway. Archives of toxicology 92(11):3347–3358 10.1007/s00204-018-2299-4

Kumari A, Tirkey NN (2019) Tenuazonic Acid: A Potent Mycotoxin. In: Singh K, Srivastava N (eds) Recent Trends in Human and Animal Mycology. Springer Singapore, Singapore, p 203–211 10.1007/978-981-13-9435-5_8

Kusunoki N, Takara K, Tanigawara Y, et al. (1998) Inhibitory effects of a cyclosporin derivative, SDZ PSC 833, on transport of doxorubicin and vinblastine via human P-glycoprotein. Jpn J Cancer Res 89(11):1220–8 10.1111/j.1349-7006.1998.tb00518.x

Li X, Jin F, Lee HJ, Lee CJ (2021) Kaempferol Regulates the Expression of Airway MUC5AC Mucin Gene via IκBα-NF-κB p65 and p38-p44/42-Sp1 Signaling Pathways. Biomol Ther (Seoul) 29(3):303–310 10.4062/biomolther.2020.149

Liu Y, Zou Z-B, Gu F-D, et al. (2024) Marine-Derived Alternariol Monomethyl Ether Alleviates Ovalbumin-Induced Food Allergy by Suppressing MAPK and NF-κB Signaling Pathways of Mast Cells. Journal of Agricultural and Food Chemistry 72(10):5463–5476 10.1021/acs.jafc.3c08126

Louro H, Vettorazzi A, Lopez de Cerain A, et al. (2024) Hazard characterization of Alternaria toxins to identify data gaps and improve risk assessment for human health. Archives of toxicology 98(2):425–469 10.1007/s00204-023-03636-8

Marin DE, Bulgaru VC, Pertea A, Grosu IA, Pistol GC, Taranu I (2024) Alternariol Monomethyl-Ether Induces Toxicity via Cell Death and Oxidative Stress in Swine Intestinal Epithelial Cells. Toxins 16(5) 10.3390/toxins16050223

Markus J, Landry T, Stevens Z, et al. (2021) Human small intestinal organotypic culture model for drug permeation, inflammation, and toxicity assays. In Vitro Cell Dev Biol Anim 57(2):160–173 10.1007/s11626-020-00526-6

Maubon N, Le Vee M, Fossati L, et al. (2007) Analysis of drug transporter expression in human intestinal Caco-2 cells by real-time PCR. Fundam Clin Pharmacol 21(6):659–63 10.1111/j.1472-8206.2007.00550.x

Mikula H, Skrinjar P, Sohr B, Ellmer D, Hametner C, Fröhlich J (2013) Total synthesis of masked Alternaria mycotoxins—sulfates and glucosides of alternariol (AOH) and alternariol-9-methyl ether (AME). Tetrahedron 69(48):10322–10330 10.1016/j.tet.2013.10.008

Olli KE, Rapp C, O’Connell L, et al. (2020) Muc5ac Expression Protects the Colonic Barrier in Experimental Colitis. Inflammatory Bowel Diseases 26(9):1353–1367 10.1093/ibd/izaa064

Partsch V, Crudo F, Piller D, Varga E, Del Favero G, Marko D (2025) Resolving complexity: Identification of altersetin and toxin mixtures responsible for the immunomodulatory, antiestrogenic and genotoxic potential of a complex Alternaria mycotoxin extract. Food and chemical toxicology : an international journal published for the British Industrial Biological Research Association 198:115315 10.1016/j.fct.2025.115315

Partsch V, Selimagić A, Crudo F, Marko D (2026) Immunomodulatory potential of Alternaria mycotoxins on immune and intestinal cells: a comparative systematic in vitro study. Archives of toxicology 10.1007/s00204-026-04346-7

Pelaseyed T, Bergström JH, Gustafsson JK, et al. (2014) The mucus and mucins of the goblet cells and enterocytes provide the first defense line of the gastrointestinal tract and interact with the immune system. Immunol Rev 260(1):8–20 10.1111/imr.12182

Pfeiffer E, Schebb NH, Podlech J, Metzler M (2007) Novel oxidative in vitro metabolites of the mycotoxins alternariol and alternariol methyl ether. MolNutrFood Res 51(3):307–316 10.1002/mnfr.200600237

Puntscher H, Aichinger G, Grabher S, et al. (2019) Bioavailability, metabolism, and excretion of a complex Alternaria culture extract versus altertoxin II: a comparative study in rats. Archives of toxicology 93(11):3153–3167 10.1007/s00204-019-02575-7

Qiao X, Zhang J, Yang Y, et al. (2020) Development of a simple and rapid LC-MS/MS method for the simultaneous quantification of five Alternaria mycotoxins in human urine. Journal of Chromatography B 1144:122096 10.1016/j.jchromb.2020.122096

Robert H, Payros D, Pinton P, Théodorou V, Mercier-Bonin M, Oswald IP (2017) Impact of mycotoxins on the intestine: are mucus and microbiota new targets? Journal of Toxicology and Environmental Health, Part B 20(5):249–275 10.1080/10937404.2017.1326071

Schmutz C, Cenk E, Marko D (2019) The Alternaria Mycotoxin Alternariol Triggers the Immune Response of IL-1β-stimulated, Differentiated Caco-2 Cells. Mol Nutr Food Res 63(20):e1900341 10.1002/mnfr.201900341

Schrader TJ, Cherry W, Soper K, Langlois I (2006) Further examination of the effects of nitrosylation on Alternaria alternata mycotoxin mutagenicity in vitro. Mutation research 606(1-2):61–71 10.1016/j.mrgentox.2006.02.008

Schrader TJ, Cherry W, Soper K, Langlois I, Vijay HM (2001) Examination of Alternaria alternata mutagenicity and effects of nitrosylation using the Ames Salmonella test. TeratogCarcinogMutagen 21(4):261–274 10.1002/tcm.1014

Sdogati S, Pecorelli I, Orsini S, et al. (2026) Alternaria toxins in tomato products: from analytical advances to risk assessment and regulatory needs. Current Research in Food Science 13:101483 10.1016/j.crfs.2026.101483

Shrivastava A, Gupta V (2011) Methods for the determination of limit of detection and limit of quantitation of the analytical methods. Chronicles of Young Scientists 2(1):4 10.4103/2229-5186.79345

Slattery O, Dahle MK, Sundaram AYM, Nowak BF, Gjessing MC, Solhaug A (2023) Functional and molecular characterization of the Atlantic salmon gill epithelium cell line ASG-10; a tool for in vitro gill research. Front Mol Biosci 10:1242879 10.3389/fmolb.2023.1242879

Snapkow I, Smith NM, Arnesdotter E, et al. (2024) New approach methodologies to enhance human health risk assessment of immunotoxic properties of chemicals — a PARC (Partnership for the Assessment of Risk from Chemicals) project. Frontiers in Toxicology Volume 6–2024 10.3389/ftox.2024.1339104

Solhaug A, Dowd GC, Dayeh VR, Sindre H, Lee LEJ, Bols NC (2025) Improve your success with fish cell lines-small things that matter. In Vitro Cell Dev Biol Anim 10.1007/s11626-025-01042-1

Solhaug A, Eriksen GS, Holme JA (2016a) Mechanisms of Action and Toxicity of the Mycotoxin Alternariol: A Review. Basic Clin Pharmacol Toxicol 10.1111/bcpt.12635

Solhaug A, Holme JA, Haglund K, et al. (2013) Alternariol induces abnormal nuclear morphology and cell cycle arrest in murine RAW 264.7 macrophages. Toxicol Lett 219(1):8–17 10.1016/j.toxlet.2013.02.012

Solhaug A, Karlsoen LM, Holme JA, Kristoffersen AB, Eriksen GS (2016b) Immunomodulatory effects of individual and combined mycotoxins in the THP-1 cell line. Toxicology in vitro : an international journal published in association with BIBRA 36:120–132 10.1016/j.tiv.2016.07.012

Solhaug A, Vines LL, Ivanova L, et al. (2012) Mechanisms involved in alternariol-induced cell cycle arrest. Mutation research 738–739:1–11 10.1016/j.mrfmmm.2012.09.001

Solhaug A, Wisbech C, Christoffersen TE, et al. (2015) The mycotoxin alternariol induces DNA damage and modify macrophage phenotype and inflammatory responses. Toxicol Lett 239(1):9–21 10.1016/j.toxlet.2015.08.1107

Tiberghien F, Loor F (1996) Ranking of P-glycoprotein substrates and inhibitors by a calcein-AM fluorometry screening assay. Anticancer Drugs 7(5):568–78 10.1097/00001813-199607000-00012

Tiessen C, Fehr M, Schwarz C, et al. (2013) Modulation of the cellular redox status by the Alternaria toxins alternariol and alternariol monomethyl ether. Toxicol Lett 216(1):23–30 10.1016/j.toxlet.2012.11.005

Tran VN, Viktorova J, Augustynkova K, et al. (2020) In Silico and In Vitro Studies of Mycotoxins and Their Cocktails; Their Toxicity and Its Mitigation by Silibinin Pre-Treatment. Toxins 12(3):148 10.3390/toxins12030148

Van De Walle J, Hendrickx A, Romier B, Larondelle Y, Schneider YJ (2010) Inflammatory parameters in Caco-2 cells: effect of stimuli nature, concentration, combination and cell differentiation. Toxicology in vitro : an international journal published in association with BIBRA 24(5):1441–9 10.1016/j.tiv.2010.04.002

Vazquez D, Barbacid M, Carrasco L (1974) Inhibitors of mammalian protein synthesis. Hamatol Bluttransfus 14:327–40 10.1016/0014-5793(74)80689-7

Wan L-YM, Allen KJ, Turner PC, El-Nezami H (2014) Modulation of Mucin mRNA (MUC5AC and MUC5B) Expression and Protein Production and Secretion in Caco-2/HT29-MTX Co-cultures Following Exposure to Individual and Combined Fusarium Mycotoxins. Toxicological Sciences 139(1):83–98 10.1093/toxsci/kfu019

Wan MLY, Co VA, Turner PC, Nagendra SP, El-Nezami H (2025) Deoxynivalenol modulated mucin expression and proinflammatory cytokine production, affecting susceptibility to enteroinvasive Escherichia coli infection in intestinal epithelial cells. Journal of Food Science 90(2):e70079 10.1111/1750-3841.70079

Won MK, Sohyun; Kim, Tae-Hyun. (2015) An Efficient Synthesis of Alternariol. Journal of the Korean Chemical Society 59(5):3 10.5012/jkcs.2015.59.5.471

Xiao J, Zhang Q, Gao Y-Q, Tang J-J, Zhang A-L, Gao J-M (2014) Secondary Metabolites from the Endophytic Botryosphaeria dothidea of Melia azedarach and Their Antifungal, Antibacterial, Antioxidant, and Cytotoxic Activities. Journal of Agricultural and Food Chemistry 62(16):3584–3590 10.1021/jf500054f

Zorova LD, Popkov VA, Plotnikov EY, et al. (2018) Mitochondrial membrane potential. Anal Biochem 552:50– 59 10.1016/j.ab.2017.07.009

