## Supplementary_Solhaug et al for "From cellular stress to cytokine modulation: An *in vitro* study of intestinal effects of major *Alternaria* mycotoxins"

**Table S1:** Overview of assays, toxins (exposure time and concentrations), and cell models.

| Assay and timepoints | AOH µM | AME µM | TeA µM | ATX-I µM | ALT µM | TEN µM | Model |
|---|---|---|---|---|---|---|---|
| Viability 24, 48, 72 h | 7.5, 15, 30, 60 | 5, 10, 20, 40 | 10, 20, 100, 200 | 5, 10, 20, 40 | 5, 10, 20, 40 | 5, 10, 20, 40 | [2]Undiff. Caco-2 |
| Cell cycle 24 h | 7.5, 15, 30 | 5, 10, 20 | 10, 20, 100 | 5, 10, 20 | 5, 10, 20 | 5, 10, 20 | [2]Undiff. Caco-2 |
| ROS level 2, 24 h | 30 | 20 | 100 | 20 | 20 | 20 | [2]Undiff. Caco-2 |
| P-gp interaction 1 h | 5, 10, 20, 40 | 5, 10, 20, 40 | 25, 50, 100, 200 | 5, 10, 20, 40 | 5, 10, 20, 40 | 5, 10, 20, 40 | [2]Undiff. Caco-2 |
| [1]Viability -/+ IL-1β 20 h | 0.3, 3, 30 | 0.2, 2, 20 | 2, 20, 100 | 0.2, 2, 20 | 0.2, 2, 20 | 0.2, 2, 20 | [3]Diff. Co-culture |
| [1]Barrier -/+ IL-1β 20 h | 0.3, 3, 30 | 0.2, 2, 20 | 2, 20, 100 | 0.2, 2, 20 | 0.2, 2, 20 | 0.2, 2, 20 | [3]Diff. Co-culture |
| [1]IL-8 sec. -/+ IL-1β 20 h | 0.3, 3, 30 | 0.2, 2, 20 | 2, 20, 100 | 0.2, 2, 20 | 0.2, 2, 20 | 0.2, 2, 20 | [3]Diff. Co-culture |
| [1]MUC5AC sec. -/+ IL-1β 20 h | 0.3, 3, 30 | 0.2, 2, 20 | 2, 20, 100 | 0.2, 2, 20 | 0.2, 2, 20 | 0.2, 2, 20 | [3]Diff. Co-culture |
| [1]Viability + IL-1β 20 h | 0.3, 30 | 0.2, 20 | 20, 100 | - | - | - | [4]EpiIntestinal |
| [1]Barrier + IL-1β 20 h | 0.3, 30 | 0.2, 20 | 20, 100 | - | - | - | [4]EpiIntestinal |
| [1]cytokine sec. + IL-1β 20 h | 0.3, 30 | 0.2, 20 | 20, 100 | - | - | - | [4]EpiIntestinal |
| Uptake, 5 h | 10 | 10 | 10 | - | - | - | [4]EpiIntestinal |
| Uptake (optimalisation) 0, 2, 4, 6, 24 h (AOH) 0, 5 h (AOH, AME, TeA) | 5, 10 | 10 | 10 | - | - | - | [5]Diff. Caco-2 |

[1]To mimic inflammation, the cells were pre-incubated with Alternaria toxin for 2 h, followed by the addition of IL-1β (0.5 ng/mL) and further incubated for 18 h; total incubation time 20 h. [2]Undiff. Caco-2 = undifferentiated Caco-2 cells in plastic plates, [3]Diff. Co-culture = differentiated Caco-2/HT29-MTX-E12 Co-culture in transwell inserts, [4]EpiIntestinal = differentiated tissue of primary cells in transwell inserts, [5]Diff. Caco-2 = differentiated Caco-2 in transwell inserts.

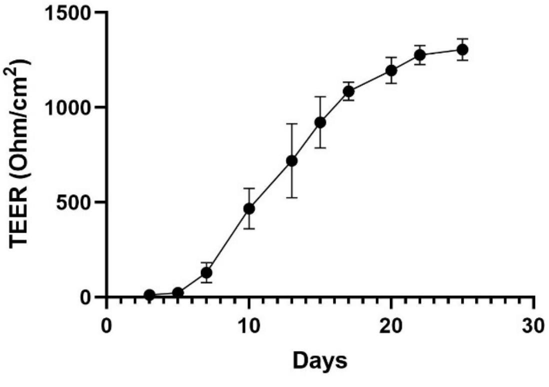


**Fig. S1:** Caco-2/HT29-MTX-E12 co-cultures grown in transwell inserts for 25 days, with regular measurements of the barrier by TEER measurements. The data represent the mean ± SD from two independent experiments done with five parallels.

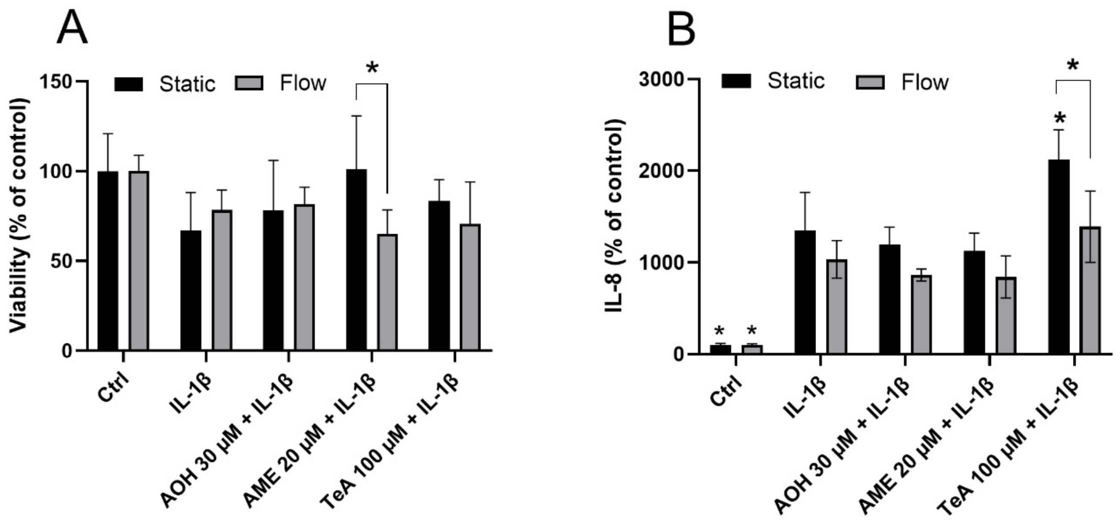


**Fig. S2:** The Caco-2/HT29-MTX-E12 co-cultures were treated with AOH 30 µM, AME 20 µM, and TeA 100 µM for 2 h before the addition of IL-1β (0.5 ng/mL) for an additional 18 h, under static or basolateral flow conditions. Viability (**A**) and IL-8 secretion (**B**) were analysed. The data are expressed as mean + SD from three independent experiments. Significant difference ($p < 0.05$) compared to IL-1β is indicated with an asterisk (*). The brackets indicate a significant difference between static and flow conditions. Two-way ANOVA with Tukey post hoc test.

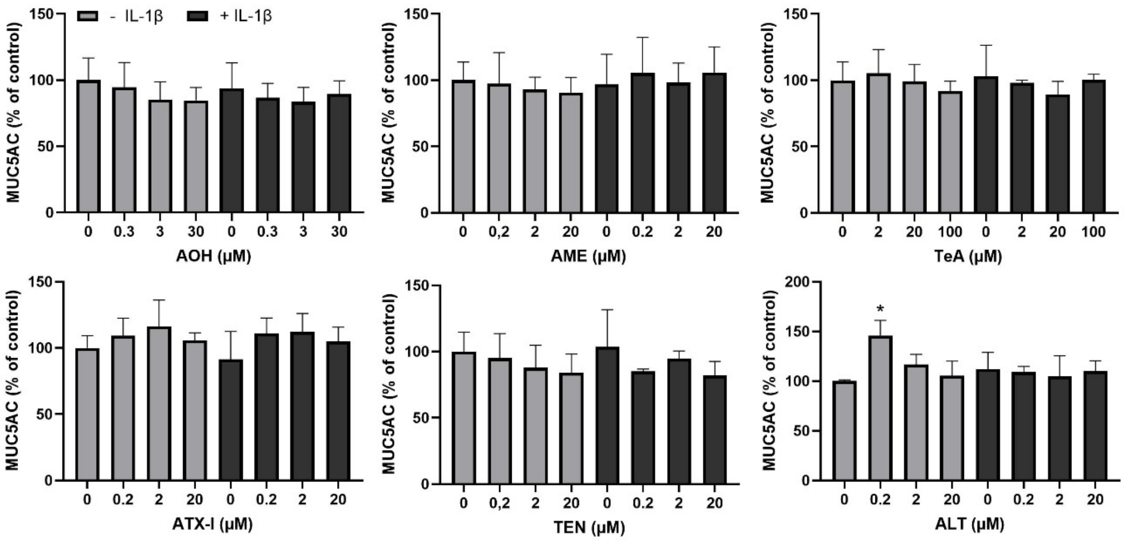


**Fig. S3:** The Caco-2/HT29-MTX-E12 co-cultures were treated with Alternaria mycotoxins as indicated for 2 h before the addition of IL-1β (0.5 ng/mL) for an additional 18 h, and MUC5AC secretions were measured by ELISA. The data are expressed as mean + SD from four to five independent experiments. Significant difference ($p < 0.05$) compared to ctrl (0 µM) is indicated with an asterisk (*). One-way ANOVA with Dunnett's post hoc test.

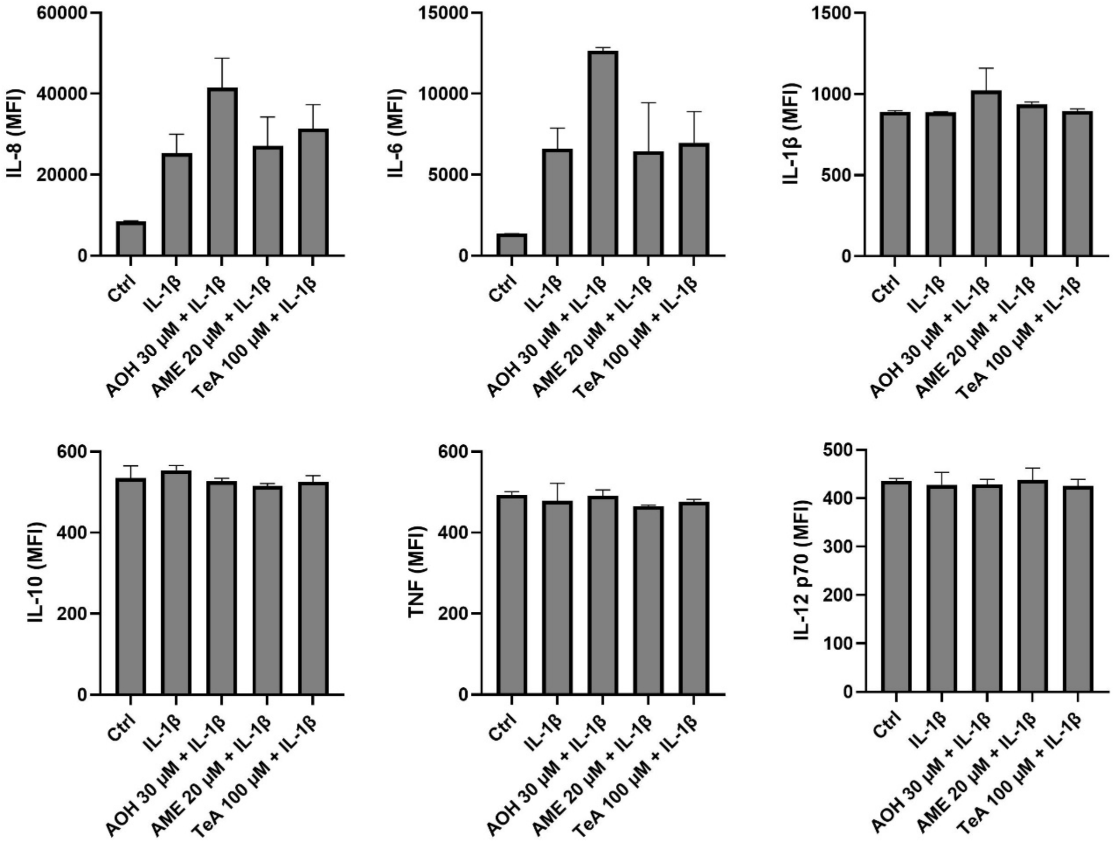


**Fig. S4:** The EpiIntestinal™ tissues were treated with AOH 30 µM, AME 20 µM, and TeA 100 µM in combination with IL-1β (0.5 ng/mL) for 20 h and simultaneously measured for IL-8, IL-6, TNFα, IL-1β, IL-10, and IL-12p70 secretion using cytometric bead array. The data are expressed as mean + SD from two independent incubations. MFI = median fluorescent intensity, relative concentration.

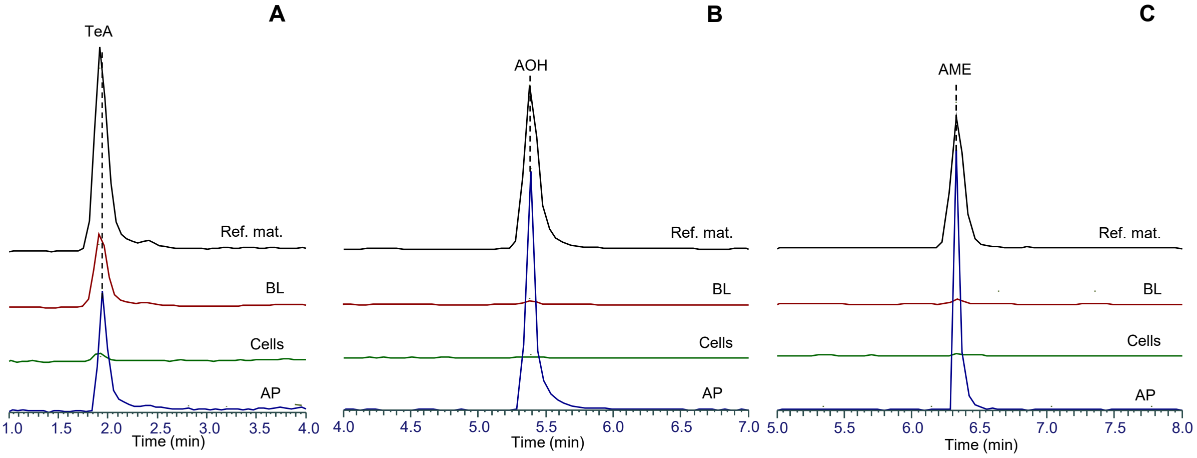


**Figure S5.** Representative extracted ion chromatograms (scaled to the most intense peak) of TeA (*m/z* 196.0979), AOH (*m/z* 257.0456), and AME (*m/z* 271.0612), detected as $[M{-}H]^-$ in apical (AP) and basolateral (BL) compartments and lysed EpiIntestinal™ tissues/cells. Verification was performed using reference material.

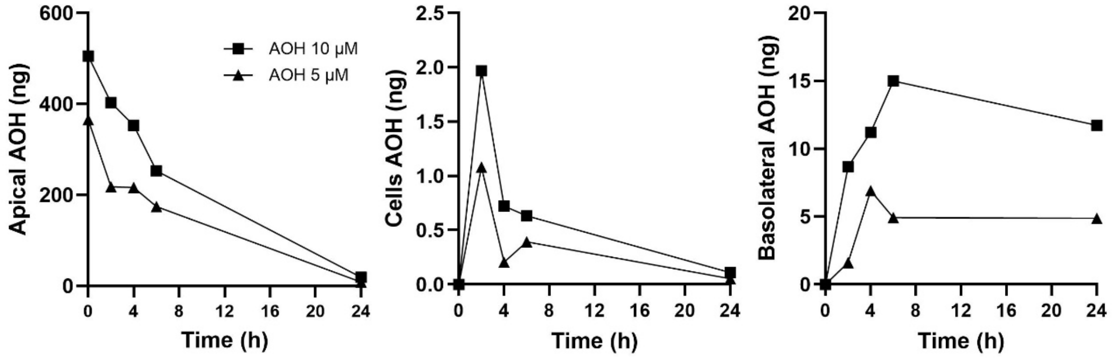


**Fig. S6:** Differentiated Caco-2 cells were apically exposed to AOH (5 and 10 µM), followed by detection of the total amount of AOH (ng) after 2 – 24 h, in the apical and basolateral compartments, and in the cells.

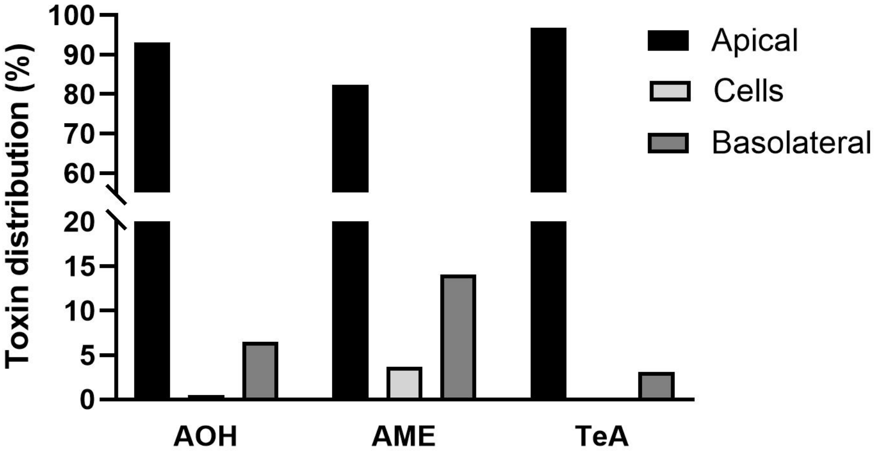


**Fig. S7**: Differentiated Caco-2 cells were apically exposed to AOH (10 µM), AME (10 µM) or TeA (10 µM) followed by LC-MS/MS analysis of AOH after 5 h, in the cell culture media (apical and basolateral) and in the cells. Also, the data analysis showed that after 5 h of exposure, the total amounts of remaining AOH, AME, and TeA are 56%, 32% and 106% of the measured starting concentration, respectively, indicating metabolism of AOH and AME.

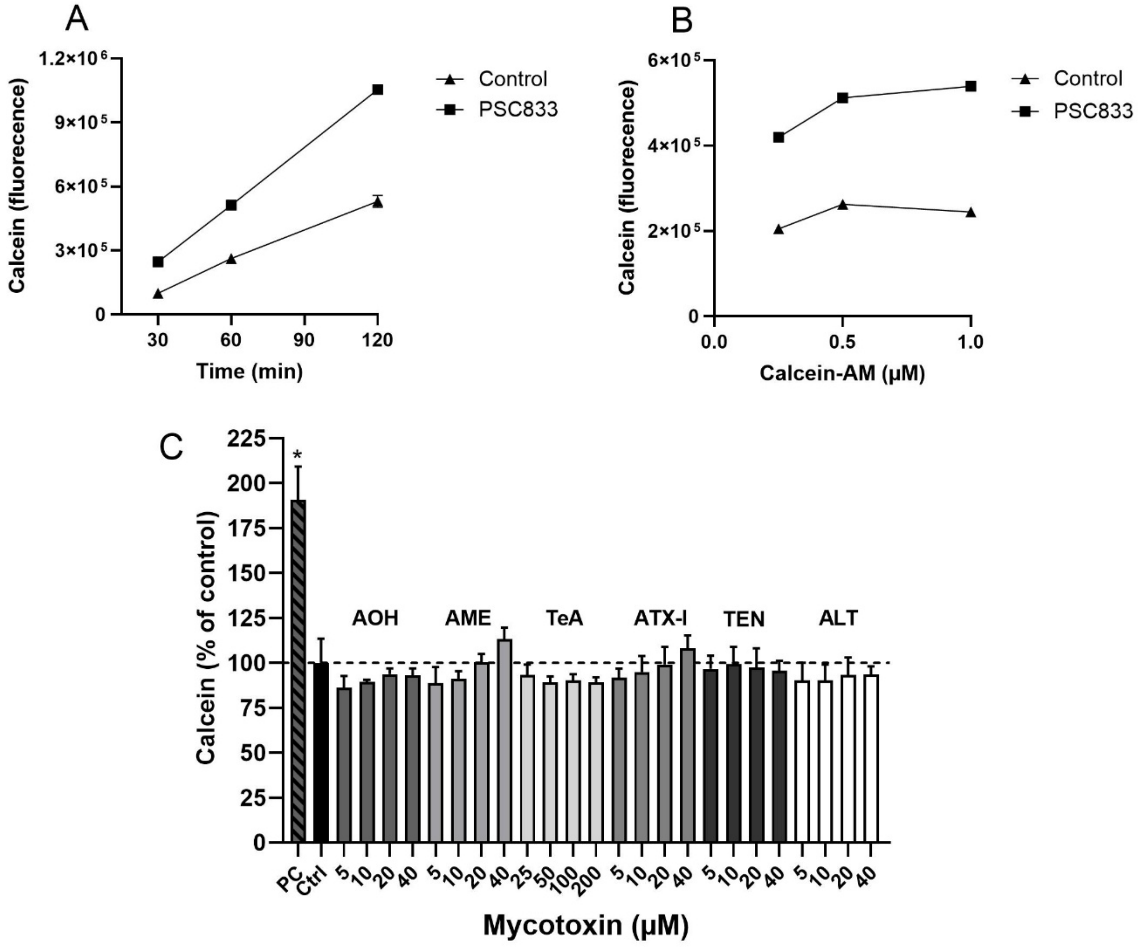


**Fig. S8:** For optimalisation of the Calcein-AM assay, undifferentiated Caco-2 cells were treated with (**A**) 0.5 µM of Calcein-AM, for different times (30, 60, 120 min) or with different concentrations of Calcein-AM (0.25, 0.5, 1.0 µM) for 1 h (**B**), or with or without the P-gp inhibitor PSC833 (5 µM). Uptake of Calcein-AM was analysed as intracellular calcein fluorescence. Undifferentiated Caco-2 cells were incubated with a wide range of concentrations of Alternaria mycotoxins, and Calcein-AM (0.5 µM) uptake was measured after 1 hr incubation (**C**). The data are expressed as mean + SD from three independent experiments, each performed in triplicate. Significant difference ($p < 0.05$) compared to control (Ctrl) is indicated with an asterisk (*). One-way ANOVA with Dunnett's post hoc test. Positive control (PC) is the P-gp inhibitor PSC833 (5 µM).
